# *Mycobacterium* trehalose polyphleates are required for infection by therapeutically useful mycobacteriophages BPs and Muddy

**DOI:** 10.1101/2023.03.14.532567

**Authors:** Katherine S. Wetzel, Morgane Illouz, Lawrence Abad, Haley G. Aull, Daniel A. Russell, Rebecca A. Garlena, Madison Cristinziano, Silke Malmsheimer, Christian Chalut, Graham F. Hatfull, Laurent Kremer

## Abstract

Mycobacteriophages are good model systems for understanding their bacterial hosts and show promise as therapeutic agents for nontuberculous mycobacterium infections. However, little is known about phage recognition of *Mycobacterium* cell surfaces, or mechanisms of phage resistance. We show here that surface-exposed trehalose polyphleates (TPPs) are required for infection of *Mycobacterium abscessus* and *Mycobacterium smegmatis* by clinically useful phages BPs and Muddy, and that TPP loss leads to defects in adsorption, infection, and confers resistance. Transposon mutagenesis indicates that TPP loss is the primary mechanism for phage resistance. Spontaneous phage resistance occurs through TPP loss, and some *M. abscessus* clinical isolates are phage-insensitive due to TPP absence. Both BPs and Muddy become TPP-independent through single amino acid substitutions in their tail spike proteins, and *M. abscessus* mutants resistant to TPP-independent phages reveal additional resistance mechanisms. Clinical use of BPs and Muddy TPP-independent mutants should preempt phage resistance caused by TPP loss.

## Introduction

Nontuberculous mycobacteria (NTM) include several important human pathogens such as *Mycobacterium abscessus* and *Mycobacterium avium*^1,2^. These infections are often refractory to effective antibiotic treatment due to both intrinsic and acquired resistance mutations, and new treatment options are needed^3^. The therapeutic application of mycobacteriophages shows some promise for the treatment of pulmonary infections in persons with cystic fibrosis^4–6^, disseminated infection following bilateral lung transplantation^4^, and disseminated *Mycobacterium chelonae* infection^7^. However, broadening therapy beyond single-patient compassionate use applications will require expansion of the repertoire of therapeutically useful phages, and increasing host range such that a higher proportion of clinical isolates can be treated^5,8,9^. Clinical administration of bacteriophages is anticipated to give rise to phage resistant mutants and disease recurrence^10^, but the frequency and mechanisms of mycobacteriophage resistance are poorly understood^11^. Very few mycobacteriophage receptors are known, although glycopeptidolipids (GPLs) are proposed as receptors for mycobacteriophage I3 in *Mycobacterium smegmatis*^12^.

Over 12,000 individual mycobacteriophages have been described, with most being isolated on *M. smegmatis*^13^. The genome sequences of 2,200 of these show them to be highly diverse genetically and pervasively mosaic^14,15^. They can be sorted into groups of genomically related phages (e.g. Cluster A, B, C etc.), some of which can be readily divided into subclusters (e.g. Subcluster A1, A2, A3 etc.) based on sequence variation^16,17^. Seven of the sequenced phages currently have no close relatives and are designated as ‘Singletons’. A subset of these phages has relatively broad host range and are also able to efficiently infect *Mycobacterium tuberculosis*, including phages in Clusters/Subclusters A2, A3, G1, K1, K2, K3, K4, and AB^18,19^. A similar subset of phages also infects some clinical isolates of *M. abscessus*, although it is noteworthy that phage host ranges on these strains is highly variable (even for related phages within clusters/subclusters), and is highly variable among different clinical isolates^4,9^. There is also substantial variation in the outcomes of phage infection of *M. abscessus* strains, with notable differences between rough and smooth colony morphotypes^9^. For example, a smaller proportion of smooth isolates are susceptible to phage infection than rough strains – as determined by plaque formation – and none of the smooth strains are efficiently killed by any phage tested^9^.

Mycobacterial cell walls characteristically have a mycolic acid-rich outer layer referred to as the mycobacterial outer membrane or mycomembrane^20^. In addition to abundant mycolic acids, there are numerous other types of complex molecules including multiple acylated lipids such as di- and polyacyltrehalose (DAT and PAT), phthiocerol dimycocerosate (PDIM) and sulfoglycolipids (SGL), although not all are found in all *Mycobacterium* species. Smooth strains of *M. abscessus* have abundant GPLs although these are lacking or greatly less abundant in rough strains^21,22^. Recently, it has been shown that some mycobacterial species, including *M. abscessus*, have trehalose polyphleates (TPPs), which are high-molecular-weight, surface-exposed glycolipids, in their cell walls^23,24^. These TPPs may be important for *M. abscessus* virulence and are associated with clumping and cording^24^. A five gene cluster, – including a polyketide synthetase (Pks) – is required for TPP biosynthesis and TPP precursor (DAT) transport to the outer surface of the cell by MmpL10^25^. TPPs are not present in *M. tuberculosis* although DAT and PAT are^25^. The specific roles of TPPs are not known, but their position on the outer surface makes them candidates for use as phage receptors.

Here, we show that TPPs are required for the binding and infection of *M. abscessus* by phages BPs and Muddy. These phages share little or no nucleotide similarity but both have been used therapeutically, sometimes in combination with each other^4,26^. *M. abscessus* transposon insertion mutants that are resistant to these phages map in all five genes involved in TPP synthesis, all have lost TPPs from their cell walls, and phage adsorption is lost. Spontaneous phage resistant mutants of some *M. abscessus* clinical isolates also have mutations in the known TPP synthesis genes, and some *M. abscessus* clinical isolates that are insensitive to BPs and Muddy are naturally defective in TPPs. However, the TPP requirement can be readily overcome by mutations in phage tail spike proteins, suggesting that TPPs are acting as a co-receptor, and the cell wall binding target of the phages is likely essential for mycobacterial viability. *M. abscessus* strains resistant to BPs and Muddy TPP-independent mutants reveal new mechanisms of phage resistance.

## Results

### Development of a transposon system for mutagenesis of *M. abscessus* clinical isolates

*M. abscessus* GD01 (subspecies *massiliense*) was selected for transposon mutagenesis as it is the first clinical isolate treated therapeutically^4^ and is killed well by phages Muddy, ZoeJΔ*45*, and BPsΔ*33*HTH_HRM10, mapping in Clusters AB, K2, and G1, respectively. Muddy is a lytic phage and ZoeJΔ*45* and BPsΔ*33*HTH_HRM10 are engineered lytic derivatives of ZoeJ^27^ and BPs^28^, respectively. Because GD01 – like many *M. abscessus* isolates – is kanamycin resistant (MIC>128 µg/mL)^29^, we re-engineered the extant Kan^R^ MycoMarT7 transposon using CRISPY-BRED^30^ to include a Hyg^R^ cassette, constructing derivatives both with and without the existing R6Kγ origin of replication (Fig. 1A). The shorter transposon (MycoMarT7-Hyg2) transduced strain GD01 ∼100 times more efficiently than the longer MycoMarT7-Hyg1 transposon; this efficiency difference was not observed for *M. smegmatis*. We transduced strain GD01 with MycoMarT7-Hyg2 and selected Hyg-resistant transductants on solid media to yield a random mutagenesis library (Fig. 1B). We note that the parent of the transposon delivery phages, TM4, does not form plaques on any *M. abscessus* strain^9^ but efficiently delivers DNA to *M. abscessus* cells^31^.

**Figure 1.**
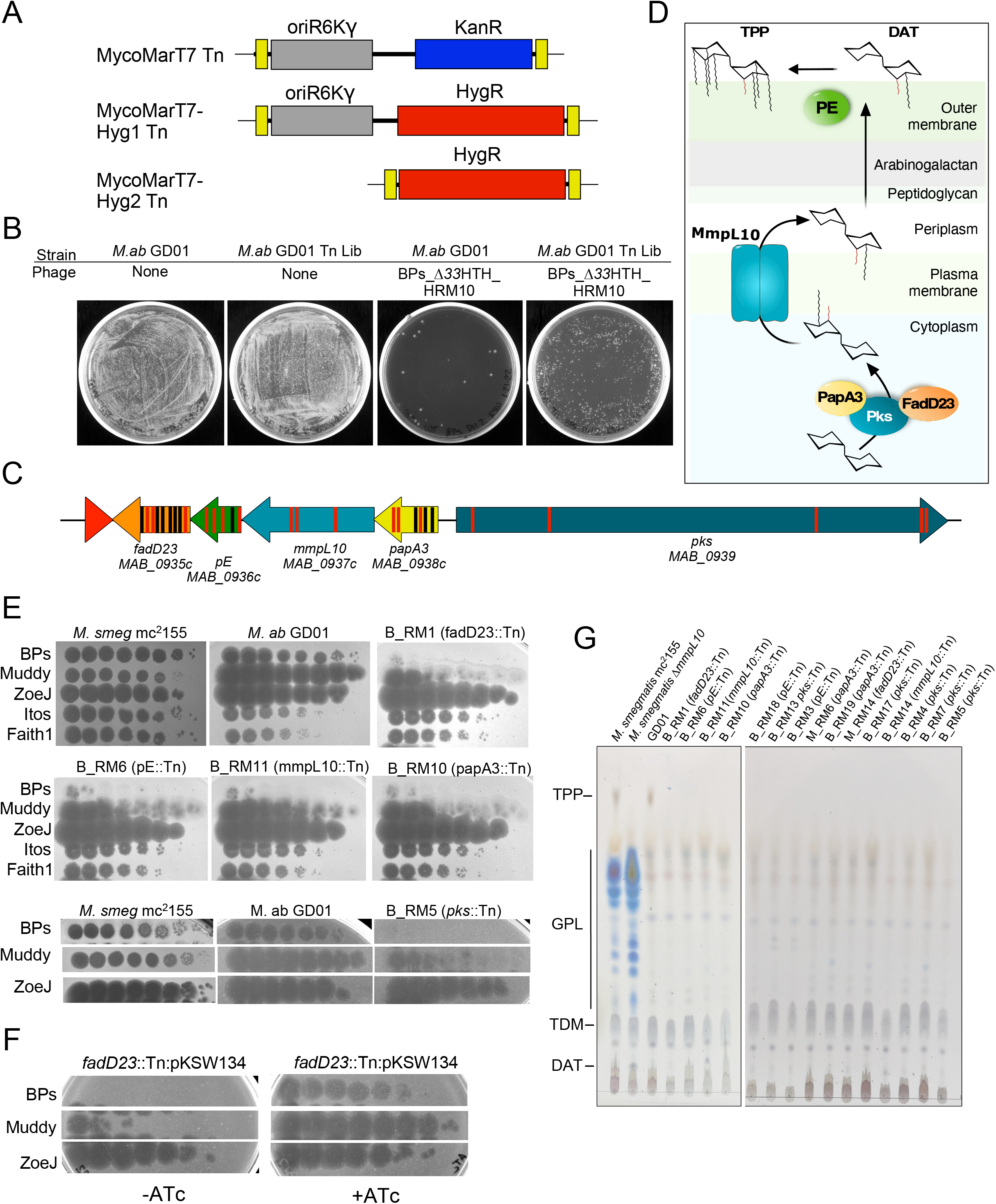
Identification of phage-resistant transposon insertion mutants of *M. abscessus* GD01. **A**. Construction of MycoMarT7-Hyg1 and MycoMarT7-Hyg2. Transposon delivery phage phiMycoMarT7 delivers a transposon containing *Escherichia coli* ori6Kγ (grey) and kanamycin resistance cassette (blue), flanked by inverted repeats (yellow boxes). CRISPY-BRED^30^ was used to create phiMycoMarT7-Hyg1 and phiMycoMarT7-Hyg2, which deliver transposons containing oriR6K (grey box) and a hygromycin resistance cassette (red box), or only a hygromycin resistance cassette. **B**. *M. abscessus* GD01 or a transposon library of *M. abscessus* strain GD01 (*M.ab* GD01 Tn Lib) were plated on solid media or solid media seeded with phage BPs_Δ*33*HTH_HRM10 (or Muddy; not shown). **C**. Locations of transposon insertions in the TPP locus in phage resistant mutants. Red and black bars show the locations of insertion isolated as resistant to BPs_Δ*33*HTH_HRM10 and Muddy, respectively. **D**. Proposed roles of Pks, PapA3, FadD23, MmpL10 and PE in the synthesis and transport of TPPs and DAT. **E**. Tenfold serial dilutions of phages were spotted onto solid media with *M. smegmatis* mc^2^155, *M. abscessus* GD01, or representative *M. abscessus* GD01 transposon insertion mutant strains: GD01Tn_BPs_HRM10_RM1 (B_RM1); GD01Tn_BPs_HRM10_RM6 (B_RM6); GD01Tn_BPs_HRM10_RM11 (B_RM11); GD01Tn_BPs_HRM10_RM10 (B_RM 10); GD01Tn_BPs_HRM10_RM5 (B_RM 5). The locations of Tn insertions are indicated in parentheses. Phages used are: BPs_Δ*33*HTH_HRM10 (“BPs”), Muddy, ZoeJΔ*43-45* (“ZoeJ”), Itos, and Faith1Δ*38-40* (“Faith1”). Plaque assays were performed at least twice with similar results. **F**. Tenfold serial dilutions of phages were spotted onto solid media with strain GD01 *fadD23*::Tn (GD01Tn_BPs_HRM10_RM1) containing plasmid pKSW134 with gene *fadD23* under expression of an anhydrotetracycline (ATc)-inducible promoter. FadD23 is not expressed in the absence of ATc (left panel) but is induced by ATc (right panel). Plaque assays were performed at least twice with similar results. **G**. TLC analysis of total lipids extracted from *M. abscessus* GD01 and mutants with transposon insertions in the TPP synthesis and transport genes. *M. smegmatis* mc^2^155 and a Δ*mmpL10* mutant strain of *M. smegmatis* are also included as controls.

### BPsΔ*33*HTH_HRM10 and Muddy Tn resistant mutants are defective in trehalose polyphleates

To identify *M. abscessus* GD01 phage resistant mutants, the Tn library was plated on solid media seeded with either BPsΔ*33*HTH_HRM10 or Muddy. Single colonies were recovered at a frequency of ∼10^-3^ and twenty individual colonies were picked from each selection, rescreened, and characterized (Fig. 1, Fig. S1, Table S1). Eighteen of the 20 BPsΔ*33*HTH_HRM10 resistant candidates were mapped, all of which have transposon insertions in a gene cluster involved in trehalose polyphleates synthesis; some appear have secondary transposon insertions mapping elsewhere (Table S1, Fig. 1C, D). Thirteen of the 20 Muddy resistant candidates were mapped and surprisingly all also contain insertions in TPP synthesis genes (Table S1, Fig. 1C, D). TPP synthesis has previously been reported to be non-essential^32^, and these observations suggest that loss of TPPs is the primary mechanism of resistance to both BPsΔ*33*HTH_HRM10 and Muddy. Further analysis showed that all of the mutants tested have similar phenotypes with a large reduction in the efficiency of plaquing (EOP) of BPsΔ*33*HTH_HRM10, and a more modest reduction in EOP of Muddy, but with formation of very turbid plaques (Fig. 1E).

Complementation of a f*adD23* Tn mutant confirmed that phage resistance results from TPP loss (Fig. 1F). All of the strains that we tested remain sensitive to ZoeJΔ*43-45*, Itos, and Faith1Δ*38-40* (Fig. 1E, S1).

Analysis of cell wall lipids shows that all of the mutants tested have lost TPPs (Fig. 1G). Interruption of TPP precursor transport (as in an *mmpL10* mutant; Fig. 1D), or loss of PE which is needed for the final step of TPP synthesis (Fig. 1D) can result in accumulation of the DAT precursor^23,25^, and our *mmpL10* transposon insertion mutants do accumulate DAT. Our *pE* mutants did not accumulate DAT and the Tn insertions may be polar, interrupting *fadD23* expression and DAT synthesis (Figs. 1C, 1D, 1G). No defects in trehalose dimycolate (TDM) synthesis were observed, which is transported by MmpL3^33^ (Fig. 1G).

### BPsΔ*33*HTH_HRM10 mutants in the predicted tail spike gene are TPP-independent

Although BPsΔ*33*HTH_HRM10 does not efficiently infect *M. abscessus* TPP synthesis mutants, plaques were observed at high phage titers that are candidates for TPP-independent mutants (Fig. 1E). Five individual plaques were purified, shown to have heritable infection of *M. abscessus* TPP mutants, and were further characterized. Two were isolated on *M. abscessus* GD01 *fadD23::Tn* (phKSW2 and phKSW3), two on GD01 *pE::Tn* (phKSW4 and phKSW5), and one on GD180_RM2 (BPs_REM1; see below); an additional mutant (phKSW1) was isolated on *M. smegmatis ΔMSMEG_5439* (Table S2; see below). These mutants form clear plaques on all TPP synthesis pathway mutants tested (Fig. 2A), and sequencing showed that all have single amino acid substitutions in the predicted BPs tail spike protein, gp22 (Table S2). Interestingly, two of these substitutions, gp22 A306V and A604E (present in phKSW3 and phKSW5, respectively), were reported previously as BPs host range mutants able to infect *M. tuberculosis*^18,28^. The gp22 A604E substitution is also present in phage BPsΔ*33*HTH_HRM^GD03^ that infects some other *M. abscessus* strains^4^. Although phKSW4 (and phKSW2; Table S2) has a gp22 L462R substitution, BPs_REM1 has both a gp22 L462R substitution and a G780R substitution. BPs_REM1 forms somewhat clearer plaques than phKSW4 on the TPP mutants (Fig. 2A) suggesting the G780R has an additive effect towards clear plaque formation.

**Figure 2.**
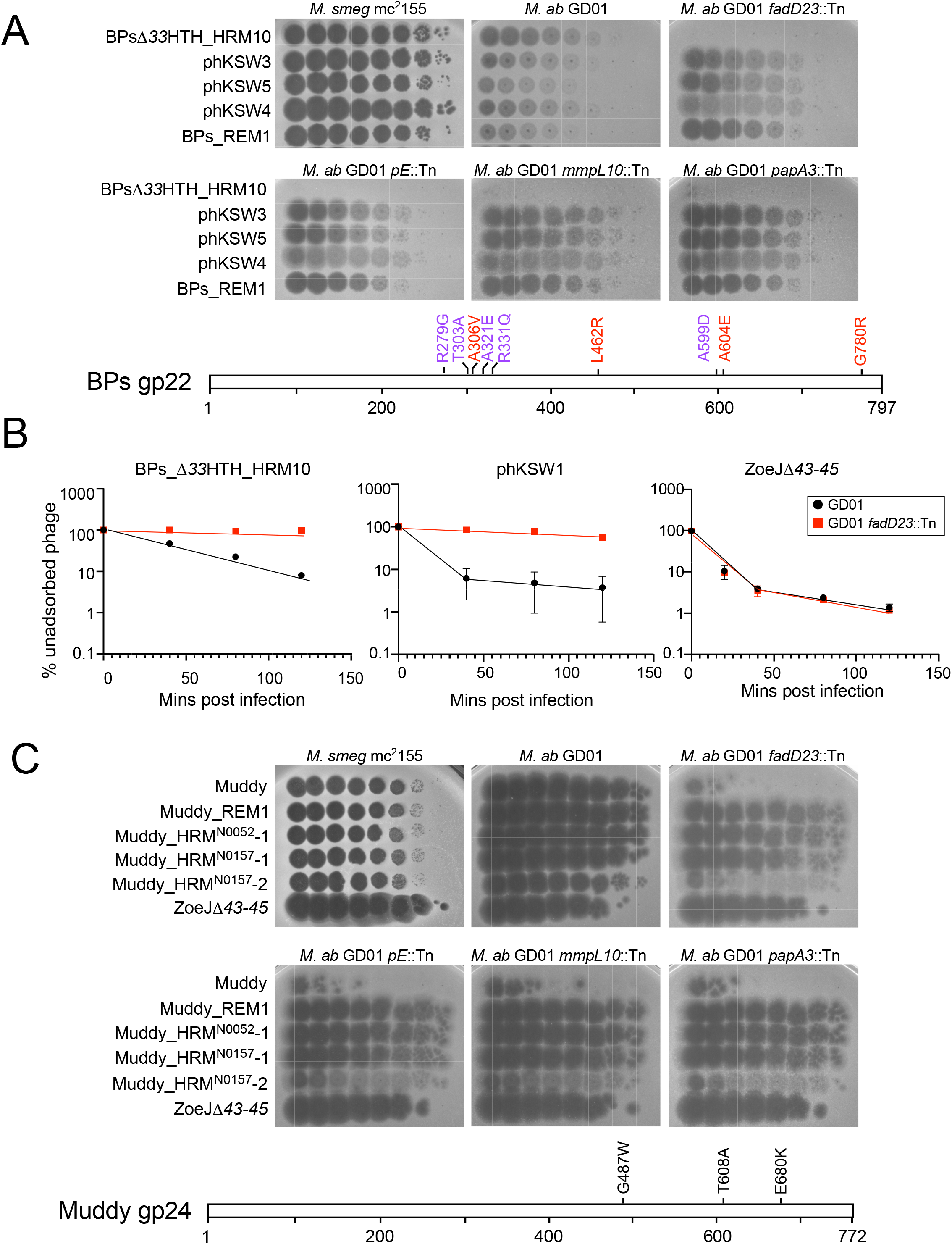
Mutants of BPs_Δ*33*HTH_HRM10 and Muddy overcome TPP loss. **A**. Ten-fold serial dilutions of BPs_Δ*33*HTH_HRM10 and gp22 mutants (as indicated on left; see Table S2) were spotted onto solid media with *M. smegmatis* mc^2^155, *M. abscessus* GD01, or *M. abscessus* GD01 transposon insertion mutant strains. Plaque assays were performed at least twice with similar results. The locations of amino acid substitutions in BPs_Δ*33*HTH_HRM10 gp22 conferring the ability to infect TPP-deficient strains (red) or were previously found to broaden host range to include *M. tuberculosis* (purple) (bottom panel). The A306V and A604E substitutions were identified with both assays. **B**. Adsorption of phages BPs_Δ*33*HTH_HRM10, phKSW1 and ZoeJΔ*43-45* to *M. abscessus* strains GD01 and GD01 *fadD23*::Tn (GD01Tn_BPs_HRM10_RM1) as indicated by the percentage of unadsorbed phages remaining in infection supernatants at different times after infection. Assays were performed in duplicate twice and the mean is displayed; error bars represent one standard deviation. **C**. Tenfold serial dilutions of Muddy and Muddy gp24 mutants (as indicated on left) were spotted onto solid media with *M. smegmatis* mc^2^155, *M. abscessus* GD01, or *M. abscessus* GD01 transposon insertion mutant strains. Plaque assays were performed at least twice with similar results. The location of amino acid substitutions in Muddy gp24 that confer the ability to infect TPP deficient strains (bottom panel).

### TPPs are required for efficient adsorption of BPsΔ*33*HTH_HRM10 to *M. abscessus* GD01

Because TPPs are surface-exposed and are required for BPsΔ*33*HTH_HRM10 infection, we tested if they are required for adsorption (Fig. 2B). Wild-type BPs adsorbs relatively poorly to *M. smegmatis*^18^ and BPsΔ*33*HTH_HRM10 adsorption is similarly poor on *M. abscessus* GD01 (Fig. 2B). However, BPsΔ*33*HTH_HRM10 is clearly defective in adsorption to a GD01 *fadD23*::Tn mutant (Fig. 2B, left panel). Interestingly, the TPP-independent phage phKSW1 (Table S2) adsorbs considerably faster to GD01 (as does a BPs gp22 A604E mutant in *M. smegmatis*^18^) than its parent phage (Fig. 2B, middle panel), and shows only a small improvement in adsorption on the GD01 *fadD23*::Tn mutant relative to BPsΔ*33*HTH_HRM10 infection of GD01 (Fig. 2B). In contrast, ZoeJΔ*43*-*45* adsorbs similarly to both *M. abscessus* strains (Fig. 2B, right panel).

### Muddy mutants in the predicted tail spike gene *24* are also TPP-independent

We similarly isolated a resistance escape mutant of Muddy (Muddy_REM1, Table 2) and – together with three Muddy mutants with expanded *M. tuberculosis*^19^ host ranges – characterized their infection of TPP pathway mutants (Fig. 2C). Three of the mutants (Muddy_REM1, Muddy_HRM^N0157^–1 and Muddy_HRM^N0052^–1) efficiently infect all of the TPP pathway mutants; Muddy_HRM^N0157^-2 forms very turbid plaques on all of the mutants similar to wild-type Muddy (Fig. 2C). Sequencing showed that Muddy_REM1 contains a single base substitution in the tail spike gene *24* conferring a E680K substitution (Table S2), the same substitution as in Muddy_HRM^N0052^-1; Muddy_HRM^N0157^-1 and Muddy_HRM^N0157^-2 have G487W and T608A substitutions in gp24, respectively^19^.

### Spontaneous *M. abscessus* phage resistant mutants in the TPP pathway

We previously reported *M. abscessus* mutants spontaneously resistant to BPs derivatives^9^. Two of the strains (GD17_RM1 and GD22_RM4, Table S3) have mutations in *pks* and are at least partially resistant to BPsΔ*33*HTH_HRM10^9^. We have similarly isolated three additional spontaneous mutants resistant to BPsΔ*33*HTH_HRM10, two of which (GD38_RM2 and GD59_RM1) have mutations in *pks*; the third (GD180_RM2) has a nonsense mutation in *mmpL10* (Table S3). BPsΔ*33*HTH_HRM10 does not form plaques on mutants GD38_ RM2, GD17_RM1 or GD59_RM1 and very small plaques at a reduced EOP on GD22_RM4 (Fig. 3A). Thus, point mutations in *M. abscessus* TPP synthesis genes can give rise to BPs resistance, although these have not been observed clinically^5^. These mutants are infected well by other phages we tested that infect the parent strain (Fig. 3A). The *M. abscessus* Pks protein (MAB_0939) is a 3697-residue multidomain protein (Fig. 3B). Two of the spontaneously resistant mutants have frameshift mutations close to the midpoint of the gene (at codons 2,115 and 2,389, Table S3, Fig. 3B) and two others have amino acid substitutions in the N-terminal Keto-synthase (KS) domain (Table S3, Fig. 3B). We note that the two frameshift mutations are in the second Acyltransferase (AT) domain and leave the upstream domains intact (Fig. 3B).

**Figure 3.**
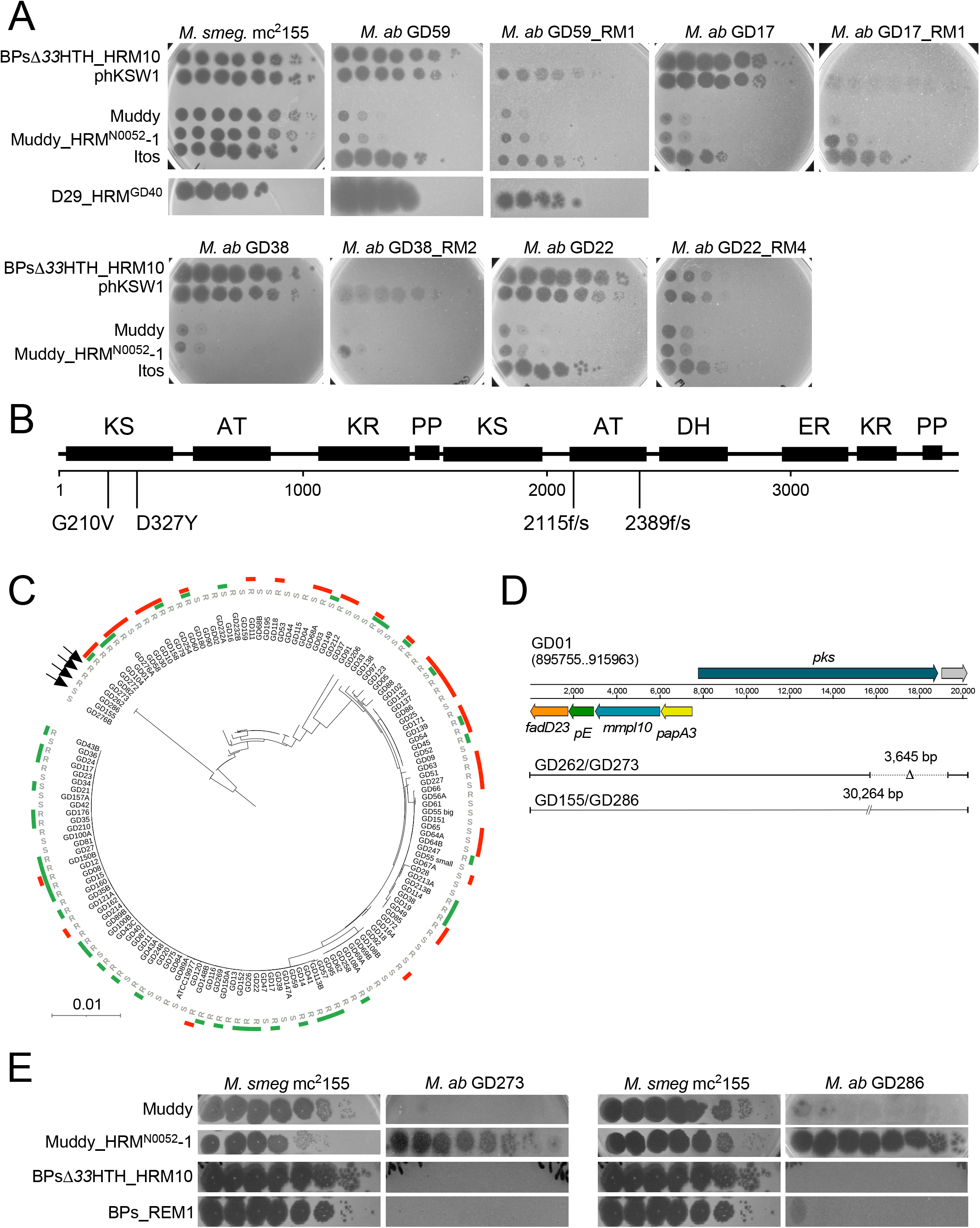
Phage infection profiles of *M. abscessus* phage resistant mutants. Tenfold serial dilutions of phage lysates (as indicated on left) were spotted onto solid media with *M. smegmatis* mc^2^155, the parent *M. abscessus* strains or spontaneously isolated phage resistant mutant (RM) derivatives. Plaque assays were performed at least twice with similar results. **B**. A schematic representation of *M. abscessus* Pks showing the location of predicted functional domains and the amino acid changes in spontaneous phage resistant mutants below. Domain abbreviations are: AT, Acyltransferase; KS, Ketosynthase, KR, Ketoreductase; DH, Dehydratase; ER, Enoylreductase; PP, phosphopantetheinylate acyl carrier protein. Domains were identified using the PKS analysis web site at http://nrps.igs.umaryland.edu/ ^50^. **C**. Amino acid sequences from the five TPP synthesis pathway genes in 143 *M. abscessus* clinical isolates (and *M. abscessus* ATCC19977) were concatenated and used to construct a phylogenetic tree. Strain morphotypes are labeled as either rough (R) or smooth (S). Susceptibilities to phages BPs_Δ*33*HTH_HRM10 and Muddy are represented in green and red, respectively^9^. Arrows indicate strains in panels D and E. **D**. Position of large deletions (GD262 and GD273) or insertions (GD155 and GD286) in the *pks* gene with respect to the GD01 TPP locus. **E**. Ten-fold serial dilutions of phage lysates (as indicated on left) were spotted onto solid media with either *M. smegmatis* mc^2^155 or *M. abscessus* strains GD273 and GD286. Plaque assays were performed at least twice with similar results.

### Some phage-insensitive *M. abscessus* clinical isolates are naturally defective in TPPs

*M. abscessus* clinical isolates vary greatly in their sensitivity to BPs_Δ*33*HTH_HRM10 and Muddy^9^. There are likely numerous determining factors, but these could include loss of TPPs. Analysis of the TPP synthesis proteins (Pks, PE, PapA3, MmpL10 and FadD23) of 143 sequenced clinical isolates and reference strain ATCC 19977 identified 37 distinct genotypes that generally correlate with global nucleotide similarity (Fig. 3C); however, no evident correlation between these variations and sensitivity to BPs_Δ*33*HTH_HRM10 and/or Muddy was observed (Fig. 3C). Most of the variations observed reflect amino acid substitutions, although two strains (GD262 and GD273) have identical large deletions in *pks* (3,645 bp) and two others (GD155 and GD286) have translocations resulting in 30.2 kbp insertions in *pks* (Fig. 3D). Both GD273 and GD286 have phage infection profiles consistent with TPP loss, and the TPP-independent mutant Muddy_HRM^N0052^–1 overcomes the defect (Fig. 3E). GD262, GD273 and GD286 are not susceptible to BPsΔ*33*HTH_HRM10 or the TPP-independent mutant BPs_REM1 (Fig. 3E), and these strains likely carry additional phage defense mechanisms targeting BPs and its derivatives. GD155 has a smooth colony morphotype (Fig. 3C) and is not susceptible to any of the phages tested here.

### Complementation restores TPP synthesis and phage infection

Mutants GD22_RM4 and GD180_RM2 – defective in *pks* and *mmpL10,* respectively (Fig. 4A) – can both be complemented to fully restore BPsΔ*33*HTH_HRM10 and Muddy infection (Fig. 4B). Both mutants lack cell wall TPPs, and TPPs are at least partially restored by complementation (Fig. 4C). Furthermore, a derivative of BPs expressing mCherry^34^ – which behaves similarly to BPsΔ*33*HTH_HRM10 in plaque assays (Fig. 4B) and liquid infections (Fig. 4D) – gives fluorescence from parent strains, but not from GD22_RM4 and GD180_RM2 (Fig. 4E, F). Complementation fully restores liquid infection of GD180_RM2, and partially restores infection of GD22_RM4 (Fig. 4D), as well as fluorescence with the reporter phage (Fig. 4E, F). These data are consistent with an early defect in phage infection in these mutants, consistent with loss of adsorption to the cell surface. We note that disruption of TPP synthesis does not interfere with Ziehl-Neelsen staining of the bacteria or alter antibiotic sensitivities (Fig. S2).

**Figure 4.**
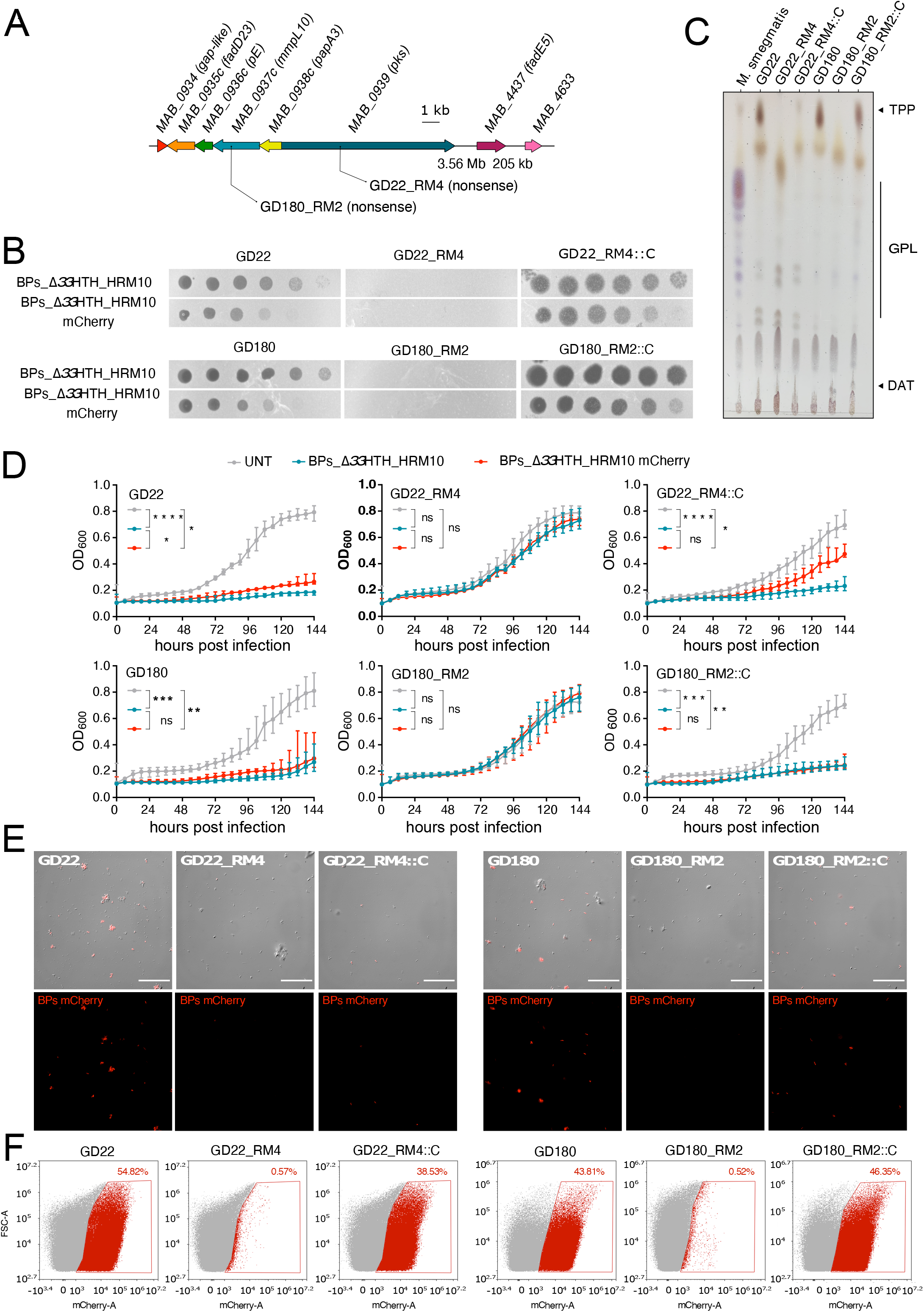
TPPs are essential for BPs_Δ*33*HTH_HRM10 to lyse *M. abscessus*. **A**. Representation of the *M. abscessus* TPP locus showing mutations affecting the clinical strains studied. **B**. Phages were spotted as ten-fold serial dilutions onto clinical strains, resistant mutants and complemented strains. Plates were incubated 2-3 days at 37° C prior to imaging. The assay was repeated at least three times and a representative experiment is shown. **C**.TLC analysis of total lipids extracted from *M. abscessus* clinical strains (GD22 and GD180), spontaneous resistant mutants (RM) and complemented strains (::C). Eluent: CHCl_3_/CH_3_OH (90:10, v/v). Anthrone was sprayed on the plates to visualize the lipid profile and followed by charring. **D**. Liquid growth of the strains with or without BPs_Δ*33*HTH_HRM10 or BPs_Δ*33*HTH_HRM10 mCherry (MOI 10) monitored every 6 hours for 6 days at 37° C in 7H9/OADC supplemented with 1 mM CaCl_2_. Data are plotted as the median of three independent experiments made in triplicates ± interquartile range. Statistical analysis conducted to compare the differences at 144h between each strain: ns, P ≥ 0.05, * P ≤ 0.05, ** P ≤ 0.01, *** P ≤ 0.001**** P ≤ 0.0001. **E**. Representative fields of *M. abscessus* clinical strains infected with BPs_Δ*33*HTH_HRM10 mCherry (MOI 10) for 4 hours at 37° C prior to fixation. Infected bacilli appear in red. **F**. Flow cytometry data are represented as dot plot show the percentage of bacilli infected with the BPs_Δ*33*HTH_HRM10 mCherry fluorophage relative to the study population.

### *M. smegmatis* mutants in the TPP synthesis pathway are resistant to phages BPs_Δ*33*HTH_HRM10 and Muddy

*M. smegmatis* is genetically tractable and susceptible to a large number of diverse phages, and using TPP mutants in *pks (MSMEG_0408)*, *mmpL10 (MSMEG_0410)*, and *pE (MSMEG_0412)*^23,35^ we showed that these have similar – albeit somewhat milder – phenotypes to *M. abscessus* TPP mutants (Fig. 5A). As expected, Δ*pks,* Δ*mmpL10* and Δ*pE* mutants failed to produce TPPs, whilst complementation restores the presence of TPPs (Fig. 5B). The relatively efficient infection of the Δ*fadD23* mutant is consistent with incomplete TPP loss, possibly due to an unidentified fatty acyl-AMP ligase partially overcoming the defect^23^. Muddy similarly forms very turbid plaques on the Δ*pks* and Δ*papA3* mutants, but only mildly so on the Δ*fadD23* mutant (Fig. 5A). Interestingly, the TPP-independent BPs and Muddy mutants infect *M. smegmatis* TPP mutants normally (Fig. 5A). Complementation of the Δ*papA3*, Δ*pks*, Δ*mmpL10*, and Δ*pE* mutants restores normal infection by both Muddy and BPsΔ*33*HTH_HRM10 (Fig. S3).

**Figure 5.**
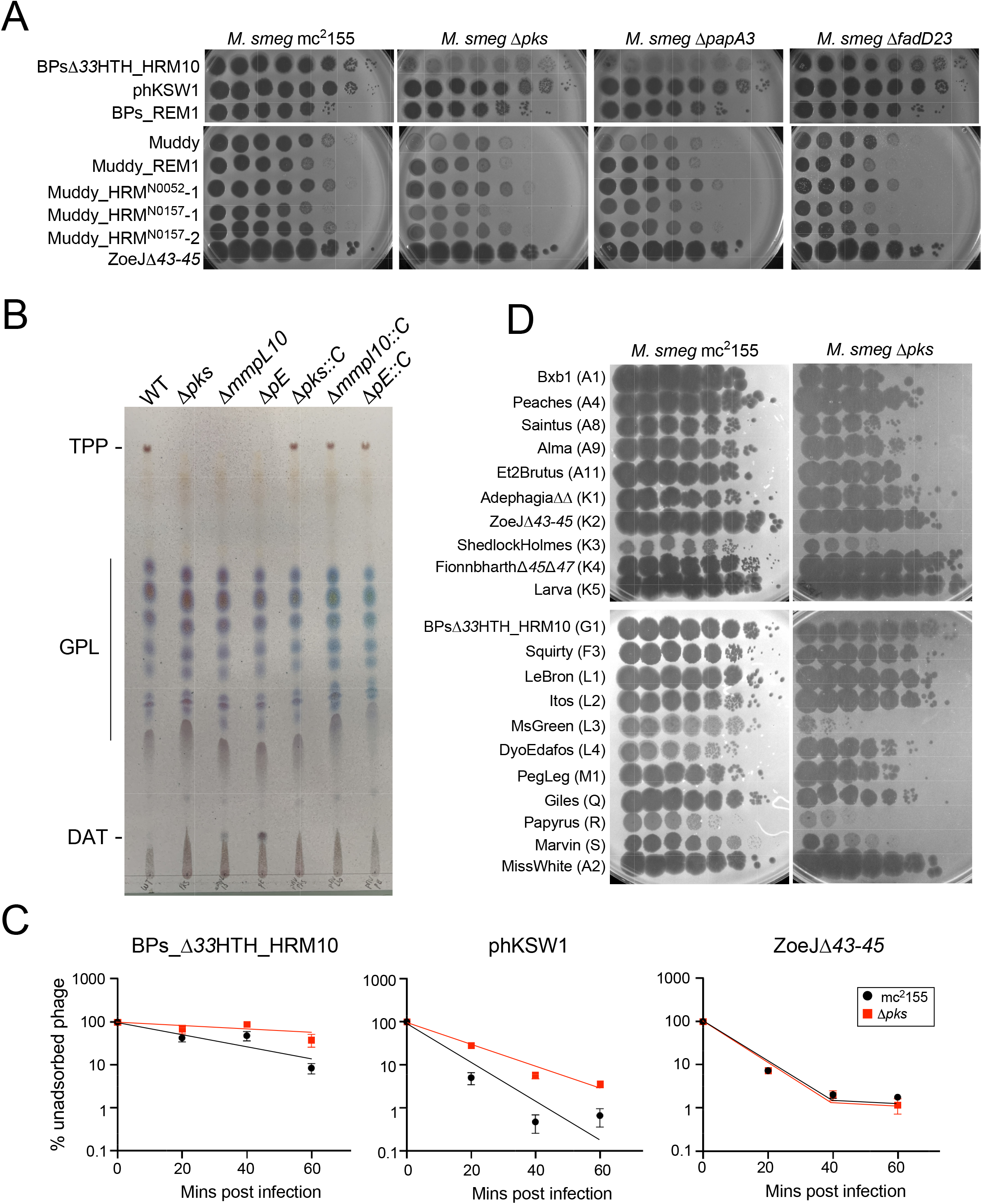
TPPs are also required for infection of *M. smegmatis* by phages BPs and Muddy. **A**. Tenfold serial dilutions of phages (as indicated on left) were spotted onto solid media with *M. smegmatis* mc^2^155, Δ*pks*, Δ*papA3* or Δ*fadD23* as indicated. Plaque assays were performed at least twice with similar results. **B**. TLC analysis of total lipids extracted from wild-type *M. smegmatis*, three TPP-deficient mutants and the corresponding complemented strains. Eluent: CHCl_3_/CH_3_OH (90:10, v/v). TLC was revealed by spraying anthrone on the plate, followed by charring. **C**. Adsorption of phages BPs_Δ*33*HTH_HRM10, phKSW1 and ZoeJΔ*43-45* on *M. smegmatis* strains mc^2^155 and Δ*pks* as indicated by the percentage of unadsorbed phages remaining in supernatant at different times after infection. Assays were performed in triplicate at least twice with similar results and a representative experiment is shown. Data are represented as the mean ±SD. Other replicates are shown in Figure S5. **D**. A panel of phages from various genetic clusters were tenfold serially diluted and spotted onto *M. smegmatis* mc^2^155 and Δ*pks.* Phage names are shown with their cluster/subcluster designation in parentheses.

BPs_Δ*33*HTH_HRM10 and its mCherry derivatives are both defective in liquid infection of the Δ*pks*, Δ*mmpL10*, and Δ*pE M. smegmatis* mutants, and efficient infection and lysis is restored by complementation (Fig. S3A, E). The mCherry reporter phage shows fluorescence in wild-type *M. smegmatis*, but loss of fluorescence in infection of all three mutants, with restoration of infection in the complemented strains (Fig. S3B, C). In addition, the mCherry fluorophage behaves similarly to its parent in plaque assays on *M. smegmatis* mutant and complemented strains (Fig. S3E). Both BPsΔ*33*HTH_HRM10 and the TPP-independent phKSW1 are defective in adsorption of a Δ*pks* mutant relative to wild type *M. smegmatis* (Fig. 5C), similar to *M. abscessus* (Fig. 2B).

Testing a broader phage panel showed that most are not dependent on TPPs, with the exceptions of ShedlockHolmes, MsGreen, and Papyrus, in Clusters/Subclusters K3, L3, and R, respectively, which show some TPP-dependence (Fig. 5D). MsGreen and ShedlockHolmes have tail genes related to Muddy gene *24*, although we note that ZoeJ does so also, and yet is not TPP-dependent. However, such variation is not unexpected, as the escape mutant observations show that only a single amino acid substitution is sufficient to confer TPP independence (Figs. 2A, 2C, 5A).

### Resistance to BPs and Muddy TPP-independent phages reveals new resistance mechanisms

The TPP-independent phage mutants infect *M. abscessus* efficiently, and we therefore repeated the selection for Tn insertion mutants to explore if there are other surface molecules required for infection. Interestingly, such mutants arise from the same library at a 100-fold lower abundance than BPsΔ*33*HTH_HRM10 and Muddy resistant mutants (Fig. 6A, Table S1). All but one of the phKSW1 resistant mutants analyzed are similarly resistant to BPsΔ*33*HTH_HRM10, BPs_REM1 and phKSW1 but remain sensitive to Muddy and Muddy_REM1 (Fig. 6B); One (RM10) is resistant to phKSW1 but is sensitive to BPs_REM1 and is resistant to Muddy but not Muddy_REM1 (Fig. 6B). One of the two Muddy_REM1 resistant mutants is only partially resistant to Muddy and Muddy_REM1, but both are fully sensitive to all of the BPs derivatives (Fig. 6B).

**Figure 6.**
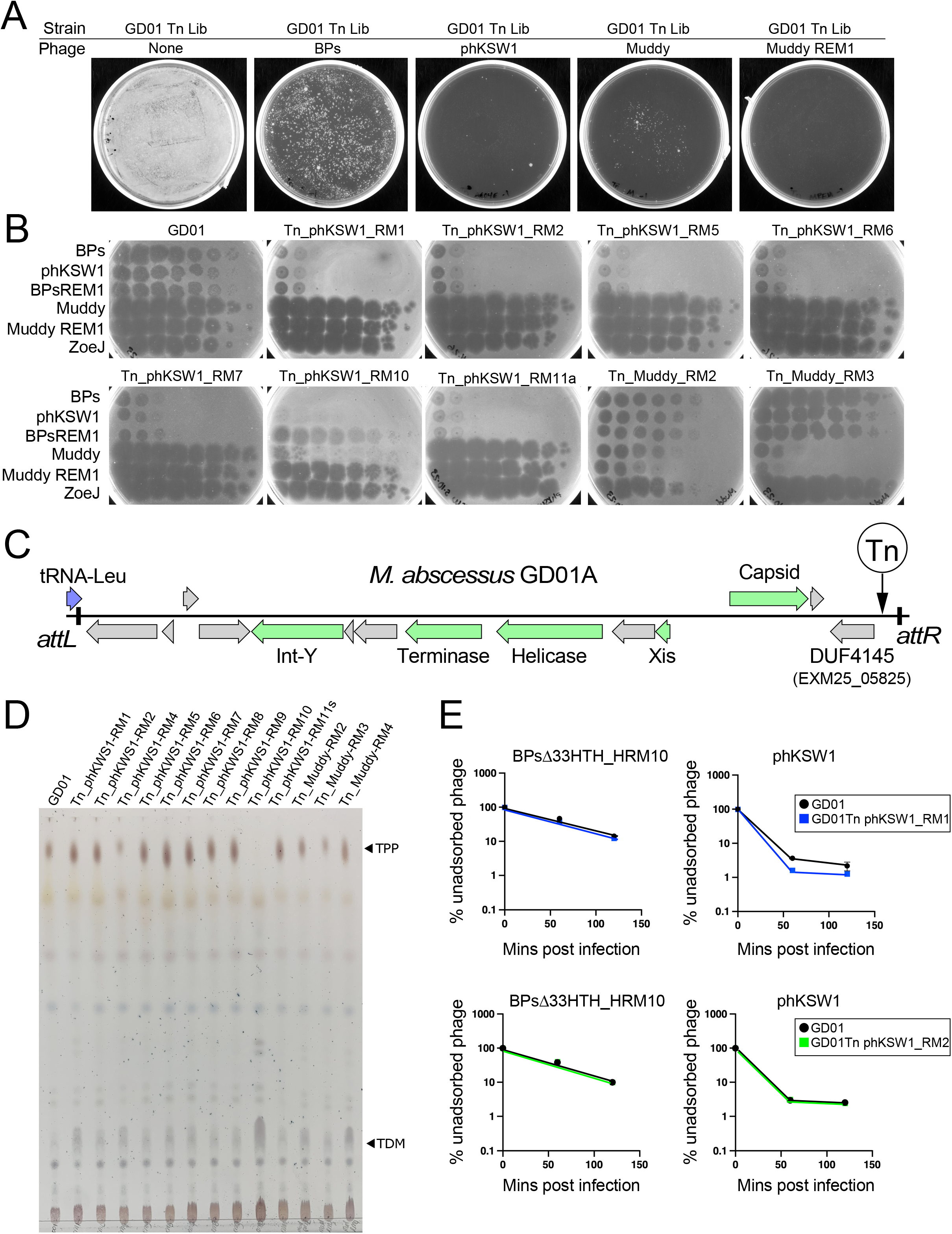
Evidence for additional phage resistance mechanisms. **A.** Recovery of *M. abscessus* GD01 transposon mutants resistant to phages BPs, phKSW1, Muddy, and Muddy_REM1. A culture of an *M. abscessus* GD01 transposon library (GD01 Tn Lib) containing approximately 10^6^ CFU was plated onto phage-seeded plates as indicated. **B**. Phage infection profiles of resistant mutants. Ten-fold serial dilutions of phages BPsΔ*33*HTH_HRM10 (“BPs”), phKSW1, Muddy, Muddy_REM1 and ZoeJΔ*43-45* (“ZoeJ”) (as indicated at the left) were spotted onto lawns of *M. abscessus* GD01 or mutants isolated as resistant to TPP-independent phages phKSW1 and Muddy_REM1. Mutant designations indicate they are Tn insertions (Tn), the phage used for mutant selection (i.e. phKSW1 or Muddy_REM1), and the mutant number (e.g. RM1, RM2 etc). Where siblings are suspected (Table S1) only one representative is shown. **C**. Organization of a putative candidate phage-inducible chromosomal island (PICI) in *M. abscessus* GD01. The satellite region is defined by the *attL* and *attR* attachment sites resulting from site-specific integration at a tRNA-Leu gene (GD01 coordinates 1158039 to 1170246). It contains several phage-related genes (green boxes) with the putative functions indicated. At the extreme right-hand end of the PICI, a gene (locus tag EXM25_05825) carries a DUF4145 domain implicated in phage defense in other systems. Four mutants contain a Tn insertion at the site indicated by the vertical arrow. **D**. TLC analysis of total lipids extracted from *M. abscessus* GD01 and transposon mutants isolated as resistant to TPP-independent phages. **E**. Adsorption of BPsΔ*33*HTH_HRM10 and phKSW1 to GD01Tn_phKSW1_RM1 (blue, top panels) and to GD01Tn_phKSW1_RM2 (green, bottom panels). Adsorption to GD01 performed in parallel is shown in black. The proportions of phage particles remaining in solution are shown at different times after infection. Assays were performed in duplicate twice and the mean is displayed; error bars represent one standard deviation.

Characterization of these mutants shows that they are unlikely to be defective in surface recognition by the phages. Two Muddy_REM1-resistant strains have Tn insertions in transcription genes *greA* and *rpoC* (Table S1), and one phKSW1 resistant mutant maps in *recB* (*MAB_0399c*; Table S1); these are unlikely to be directly involved in phage binding. Four of the phKSW1-resistant mutants (RM2, RM5, RM7, and RM8), representing at least three independent insertions (Fig. 6, S5) have transposons at GD01 coordinate 1,169,901, in a region absent from ATCC19977 and many other *M. abscessus* strains and within a candidate ‘phage-inducible chromosomal island’ (PICI) (Fig. 6C, Table S1). The insertions are upstream of GD01 gene *EXM25_05825* encoding a protein with a DUF4145 domain, which is implicated in a variety of viral defense systems and is often fused with restriction endonucleases and abortive infection systems^36–38^ (Fig. 6C). It is plausible that the BPs resistant phenotype results from overexpression of this gene.

Analysis of the cell wall lipids shows that all of the mutants retain TPPs with the exception of GD01Tn_phKSW1_RM10 (Fig. 6D), which has a Tn insertion in *papA3* in addition to a secondary insertion in *MAB_1686* (Table S1). Two additional mutants (GD01Tn_phKSW1_RM1 and RM4) have an insertion in the nearby *MAB_1690* gene and have normal TPPs (Fig. 6D); *MAB_1686* and *MAB_1690* are within a large (22 kbp) operon encoding an Mce4 transport system (Table S1). However, this Mce4 system is likely not acting as a receptor as the Tn_phMSW1_RM1 mutant does not have an adsorption defect (Fig. 6E). The Tn_phKSW1_RM2 mutant also does not have any adsorption defect (Fig. 6E).

## Discussion

Trehalose polyphleates are among the largest known lipids in mycobacteria and are structurally related to sulfolipids SL-1 and polyacylated trehalose PAT, which in contrast to TPPs, are found exclusively in *M. tuberculosis*. The roles of TPPs in mycobacterial physiology and/or growth remain unclear, but are implicated in clumping and cording in *M. abscessus*^24^. Many TPP-defective *M. abscessus* strains have rough morphotypes – typically associated with cording – consistent with the rough colony morphology primarily resulting from GPL loss^22^. Clearly, TPPs are critical for adsorption of several phages, including the therapeutically useful Muddy and BPs^4,5^. The finding that both phages require TPPs is a surprise, as they are genomically distinct, share few genes, and were thus considered to be suitable for combination in phage cocktails. Nonetheless, the availability of TPP-independent phage mutants provides substitutes to which resistance both occurs at a much lower frequency, and such mutants do not typically show co-resistance to the two phages. We propose that the TPP-independent phage replace their cognate parent phages in therapeutic cocktails.

A simple explanation for the role of TPPs is that they are specifically recognized and bound to by BPs, Muddy and the other TPP-dependent phages as the only requirement for DNA injection. However, it is then unclear as to how the TPP-independent phages overcome TPP loss, and it seems implausible that they gain the ability to bind to a completely different receptor. A more likely explanation is that TPPs act as co-receptors for Muddy and BPs and facilitate recognition of a different surface molecule; the TPP-independent phage mutants would then simply bypass the need for activation by TPPs. The observation that tail spike mutants such as phKSW1 adsorb substantially better than the parent phage, even to TPP-containing host cells, is consistent with this latter explanation. Furthermore, wild-type BPs does not efficiently infect *M. tuberculosis* H37Rv, but a mutant with the gp22 A604E substitution enables efficient infection^18^, even though *M. tuberculosis* lacks TPPs^39^. Similarly, wild-type Muddy efficiently infects *M. tuberculosis* H37Rv^19^ despite its lack of TPPs, and Muddy tail spike substitutions expand its host range to other *M. tuberculosis* strains^19^. These observations not only suggest that TPPs are not the receptors *per se* for these phages, but that there may be general mechanisms governing receptor access by the phages, together with phage strategies for expanding host cell recognition and infection. Furthermore, if TPPs are not the target of direct phage recognition, the true receptor is likely to be encoded by genes that are essential for mycobacterial viability. Thus, although transposon insertion mutagenesis has been used in other systems for identifying phage receptors^40–43^, this maybe of more limited use in *Mycobacterium*, although as we have shown here, it is useful for mapping a plethora of resistance mechanisms.

Understanding the roles of TPPs in *M. abscessus* is important for therapeutic phage use, and we note that in at least some clinical isolates the loss of TPPs through gene deletions or translocation leads to loss of infection by BPsΔ*33*HTH_HRM10 or Muddy. In the first therapeutic use of mycobacteriophages, BPsΔ*33*HTH_HRM10 and Muddy were used in combination with ZoeJ, and it is of interest that ZoeJ is not TPP-dependent. We also note that resistance to BPs derivatives or Muddy has not been observed in clinical use, even in 11 cases where only a single phage was used^5^. It is plausible that resistance through TPP loss has a tradeoff with fitness, and the roles of TPPs in *M. abscessus* pathogenicity warrant further study.

Finally, transposon mutagenesis and selection of mutants resistant to the TPP-independent phages reveals additional mechanisms of phage resistance. Particularly intriguing is the isolation of insertions in a candidate PICI, with the potential to activate expression of a PICI gene implicated in phage defense. These *Mycobacterium* PICIs and their roles in phage infection profiles deserve further investigation.

## Supporting information

All supplementary Information

## Acknowledgements

We thank Chidiebere Akusobi, Mark Sullivan, Kerry McGowen, Carlos Rodriguez and William DePas for helpful discussions. We also thank all of the students and faculty in the SEA-PHAGES program that discovered and characterized phages used in this study, and the numerous colleagues that provided *M. abscessus* strains in the GDxx series. Microscopy and flow cytometry were performed at the Montpellier Imaging Center for Microscopy (MRI). This work was supported by grants from the National Institutes of Health grants GM131729 and the Howard Hughes Medical Institute Grant GT12053 to GFH, a Cystic Fibrosis Foundation Postdoc Fellowship WETZEL21F0 to KSW, by the French National Research Agency ANR-21-CE44-0027-01 (MYCOLT) to LK and CC and by Vaincre la Mucoviscidose (RF20200502678) and the Association Grégory Lemarchal for funding the PhD fellowship of MI.

## Materials and Methods

### Bacterial strains and culture conditions

Bacterial strains (Table S4) were grown in Middlebrook 7H9 media (BD Difco) supplemented 10% oleic acid, albumin, dextrose, catalase (OADC enrichment) (7H9/OADC) or on Middlebrook 7H10/OADC solid media (BD Difco) at 37° C. Antibiotics were added when required. Transformations of electrocompetent mycobacteria were performed using a Bio-Rad Gene pulser (25 µF, 2500 V, 800 Ohms). For some *M. smegmatis* strains, Tween-80 (0.05%) was used in starter cultures but omitted in subcultures used for phage infections. Cultures used in phage infection were supplemented with 1 mM CaCl_2._ When required, *M. abscessus* strains were selected with 1 mg/mL hygromycin or 200 µg/mL streptomycin, and *M. smegmatis* were selected with 50 µg/mL hygromycin. *M. abscessus* strains in the GDxx series are part of the strain collection at the University of Pittsburgh and were kindly provided by numerous colleagues.

### Engineering of MycoMarT7-Hyg2

Phage MycoMarT7-Hyg1 and MycoMarT7-Hyg2 were engineered from phage MycoMarT7 using CRISPY-BRED recombineering^30^. Briefly, dsDNA recombineering substrates were designed that contained the desired mutation (a HygR cassette) with flanking sequences to permit the replacement of either KanR (MycoMarT7-Hyg1) or KanR and oriR6K (MycoMarT7-Hyg2). These substrates and genomic DNA from MycoMarT7 were transformed into *M. smegmatis* mc^2^155 recombineering cells that contain the plasmid pJV138^30^. Transformations were combined with cells containing a CRISPR plasmid selecting against the parent MycoMarT7 and plated on solid media; this enriches for mutants containing the allelic replacement that form plaques on the plate. Resulting plaques were screened for presence of the Hyg-marked transposon by PCR, and positive plaques were plaque purified, whole genome sequenced, and confirmed to have retained temperature sensitivity.

### Construction of the *M. abscessus* GD01 transposon insertion library and phage challenge

The transposon mutagenesis library was largely prepared as previously described^31^. Briefly, 50 mL of *M. abscessus* GD01 was grown to an OD_600_ of 0.2. The cells were pelleted and resuspended in 1 mL of phage buffer (10 mM Tris HCl, pH 7.5; 10 mM MgSO_4_; 68.5 mM NaCl; 1 mM CaCl_2_), pre-warmed to 37° C and infected with 800 µL MycoMarT7-Hyg2 (5×10^10^ pfu/mL). The cells and MycoMarT7-Hyg2 were incubated at 37° C for 7.5 hours. Cells were pelleted and resuspended in 8 mL PBS + 0.05% Tween80, and the resuspension was combined with 8 mL 40% glycerol for freezing at −80° C. The transduction frequency was determined by measuring hygromycin resistant colonies/mL and 120,000 transductants were plated onto large square plates containing solid 7H10/OADC media with 0.1% Tween80 and 1 mg/mL hygromycin. Plates incubated at 37° C for nine days. To harvest the library, cells were scraped off the solid media, resuspended in 7H9/OADC combined with 40% glycerol, aliquoted and frozen at −80° C.

To identify GD01 insertion mutants that were resistant to phage infection, ∼20 µL of GD01 Tn library was thawed and grown overnight to an OD_600_ of 0.175. Dilutions of this culture (approximately 10^4^, 10^5^ and 10^6^ cells) were spread onto 7H10/OADC solid media plates seeded with or without 10^8^ pfu of phages BPs_Δ*33*HTH_HRM10, Muddy phKSW1 or Muddy_REM1. Plates were incubated at 37° C for seven days. Colonies able to grow on phage-seeded plates were subjected to PCR to identify a transposon insertion site (see below) and struck out two times to remove any remaining phage. After streaking, single colonies were grown in liquid media and used for phage susceptibility testing by standard plaque assay.

### Identification of transposon insertion sites

Transposon insertion sites were identified by PCR using a primer that annealed to the transposon in the hygromycin resistance gene (Tn_Hyg_Fwd_2: 5’-CTTCACCTTCCTGCACGACT-3’), and a primer with a degenerate 3’ end, or if that did not yield an amplicon, nested PCR with primers Tn_Hyg_Fwd_2 and Primer 557 (5’-GGCCAGCGAGCTAACGAGCANNNNNNNGTT-3’) followed by PCR with primers Primer 414 (5’-GGCCAGCGAGCTAACGAGAC-3’) and Tn_Hyg_Fwd_1 (5’-TTCGAGGTGTTCGAGGAGAC-3’). Amplicons were gel extracted, Sanger sequenced from the transposon, and the result aligned to the GD01 sequence to identify the transposon insertion site. For most strains (and at least one strain per interrupted gene), the transposon insertion site was confirmed by designing primers that flanked the site identified by the initial PCR and confirming that this region had increased in size by 1,259 bp compared to strain GD01.

### Screening of phage susceptibility

Phage susceptibility profiles were assessed using standard plaque assays. Top agar bacterial lawns were made by combining Middlebrook Top Agar (MBTA; Middlebrook 7H9, 1 mM CaCl_2_, 0.35% BactoAgar) with 300 µL-500 µL cell culture. After top agar had solidified, phages were 10-fold serially diluted and spotted onto the top agar bacterial lawns and incubated 24-48 hours (*M. smegmatis*) or 5-7 days (*M. abscessus*) until bacterial lawns were confluent. Phages used in this study were obtained from University of Pittsburgh and *M. smegmatis* mc^2^155 was used to propagate them.

### Plasmid construction

To create plasmid pKSW131, *fadD23*, *pE*, *mmpL10* and *papA3* and the flanking intergenic sequence was amplified using Q5 HiFi 2× MasterMix (NEB) from genomic DNA isolated from *M. abscessus* GD01. The amplicon was purified and cloned into EcoRI-digested vector pLA155 using the NEBuilder Gibson Assembly Master Mix and transformed into *E. coli* strain DH5a; plasmids and primers are shown in Table S5. The culture that yielded a successfully constructed plasmid was grown at 30° C rather than 37° C, although it is unknown if this contributed to successful plasmid maintenance in the culture. To create plasmid pKSW134, the open reading frame of *fadD23* was amplified from GD01 gDNA and cloned into Pml I-digested anhydrotetracycline (ATc) inducible vector pCCK39^44^ using the NEBuilder Gibson Assembly Master Mix. The entire plasmids were sequenced by Plasmidsaurus Inc (https://www.plasmidsaurus.com/).

pMV*pks*_*mWasabi* and pMV*mmpL10*_*mWasabi* were constructed based on pMV*pks* and pMV*mmpL10* by in-fusion cloning. The *mWasabi* sequence under the control of the constitutive *Pleft\** promoter^45^ was amplified by PCR using a Q5 high-fidelity DNA polymerase (New England Biolabs). Plasmids were linearized with KpnI-HF (New England Biolabs). Agarose gels were used to purify linear fragments, then circularized using In-Fusion SNAP Assembly Master Mix (Takara Inc.), according to the manufacturer’s instructions. Stellar competent cells were used for transformation. Plasmids generated were verified by sequencing.

### Isolation and whole genome sequencing of phage resistant mycobacteria mutants

Two of these phage resistant mutants (GD17_RM1 and GD22 RM_4) were described previously^9^; the others were isolated in the same manner. Briefly, 10^8^ CFU *M. abscessus* were incubated with 10^9^ PFU of phage. Infections were plated on solid media at two and five days-post infection, and survivors were purified, tested for phage resistance, and sequenced. The mutants were sequenced as described previously^9^.

### Isolation of BPs_Δ*33*HTH_HRM10 mutants

Clear plaques were observed within high titer spots for phage BPs_Δ*33*HTH_HRM10 on strains GD01Tn_BPs_HRM10_RM6, GD01Tn_BPs_HRM10_RM11 and GD180_RM2. These plaques were picked and plated on the resistant strain two additional times to purify. A purified plaque was then used to produce a high-titer phage lysate on *M. smegmatis* mc^2^155, and subsequently subjected to gene *22* PCR sequencing or whole-genome sequencing.

### Total lipids extraction of mycobacteria and TLC analysis

Bacteria were grown in LB medium at 37° C without agitation and pelleted by centrifugation (3,000 *g*, 10 min, room temperature). Lipids were extracted as described previously^23^. Briefly, bacterial pellets were treated successively with CHCl_3_/CH_3_OH (1:2) and CHCl_3_/CH_3_OH (2:1), washed with water and dried. Lipids were resuspended in CHCl_3_ prior to spot on TLC. For TLC analysis, silica gel G60 plates (10 x 20 cm, Macherey-Nagel) were used to spot samples and lipids were separated with CHCl_3_/CH_3_OH (90:10, v/v). Lipid profiles were shown by spraying the plates with a 0.2% anthrone solution (w/v) in concentrated H_2_SO_4_ and charring.

### Adsorption Assays

*M. smegmatis* strains were grown to an OD_600_ of 0.5-0.8, then concentrated approximately tenfold to 1.75×10^9^/mL. One milliliter of cells (*M. smegmatis* mc^2^155 or mc^2^155 Δ*pks*) were infected in triplicate in a 12-well plate at a multiplicity of infection (MOI) of 0.001. At each timepoint, 50 µL of liquid were removed, pelleted, and the supernatant that contained unbound phage was titered on *M. smegmatis*. For *M. abscessus* GD01 and *M. abscessus* GD01 *fadD23*::Tn, the same protocol was followed but the strains were grown to an OD_600_ of 0.15-0.25 and cells were concentrated approximately tenfold to 6.3×10^8^/mL.

### Growth curves of mycobacteria incubated with phages

Bacterial growth assays were performed in 96-well plates (Falcon), each well containing 100 µL of bacterial culture and 100 µL of phage lysates or medium as control. Exponential phage cultures of mycobacteria were used and set at 3×10^7^ CFU/mL in Middlebrook 7H9/OADC supplemented with 1 mM CaCl_2_. Phages were incubated at a MOI 10 and diluted in 7H9/OADC supplemented with 1 mM CaCl_2_. Measurements were taken every three hours for *M. smegmatis* strains and six hours for *M. abscessus* strains using a spectrophotometer (Tecan, infinite 200 PRO) until stationary phase was reached (2 days for *M. smegmatis* strains and 6 days for *M. abscessus* strains). Plates were incubated at 37° C without agitation.

### Microscopy and flow cytometry sample preparation

Mycobacteria were sub-cultured in 7H9/OADC with agitation to obtain exponential phase cultures. Bacteria were concentrated to obtain a sample containing 1.2×10^7^ CFU (for microscopy) or 6×10^6^ CFU (for flow cytometry) and then incubated with either medium or phage BPs_Δ*33*HTH_HRM10 (MOI 10) as controls or phage BPs_Δ*33*HTH_HRM10 mCherry (MOI 10). For *M. smegmatis* strains, the infections were performed for two hours and four hours for *M. smegmatis* and *M. abscessus*, respectively, at 37° C without agitation. After infection, samples were fixed with PFA 4% for 20 min at room temperature. Samples were then diluted as necessary depending on the experiment with 7H9/OADC supplemented with 0.025% tyloxapol and sonicated to disrupt bacterial aggregates. For microscopy, samples were then mounted between coverslips and slides with Immu-Mount (Epredia). Samples were kept at 4° C in the dark until analysis.

### Microscopy

DIC (Differential interference contrast) and epifluorescence images were acquired on a ZEISS Axio Imager Z1 up-right microscope. A 63X Plan Apochromat 1.4 NA oil objective and a 100X Plan Apochromat 1.4 NA oil objective was used respectively for *M. smegmatis* or *M. abscessus* strains. mCherry was excited with an Intenslight fiber lamp with Texas Red (Ex: 560/40, dic. 585, Em: 630/75) filter cube. Images were acquired with a scMOS ZYLA 4.2 MP.

### Image analysis

Representative fields without technical artefacts were chosen. Fiji software was used to adjust intensity, brightness and contrast (identically for compared image sets).

### Flow cytometry

Infected bacteria were analysed by flow cytometry using a NovoCyte ACEA flow cytometer (excitation laser wavelength: 561 nm, emission filter: 615/20 nm). Gates were drawn using SSC-A/FSC-A and multiple cells were excluded with SSC-H / SSC-A. Uninfected cells and bacteria infected by non-fluorescent phage were included as controls. Experiments were performed at least twice with similar results. Approximately 300,000 events were recorded per experiment. Analysis was done with NovoExpress.

### Ziehl-Nilseen staining

Concentrated cultures were fixed on glass slides by heating at 150°C for 15 min followed by chemical fixation with methanol. BD Carbolfuchsin kit was used following manufacturer recommendations. Samples were observed using an Evos M7000 Imaging System.

### Drug susceptibility testing

The CLSI guidelines^46^ were followed to determine the minimum inhibitory concentrations (MICs). Briefly, all cultures were incubated in Cation-adjusted Mueller-Hinton Broth (CaMHB) at 30°C prior experiments. Each well of a 96-well plate was filled with 100 μL of bacterial suspension previously inoculated with 5.10^6^ CFU/mL, except for the first column, to which 198 μL of the bacterial suspension was added. 2 μL of drug at its highest concentration was added to the first column containing 198 μL of bacterial suspension and was two-fold serially diluted. Results are obtained after 4 days of incubation at 30°C without agitation. Three independent experiments were carried out in duplicate.

### Statistical analysis

Statistical analysis was carried out with GraphPad Prism version 9.0.0 for Windows (GraphPad Software, San Diego, California, United States). Descriptive statistics are cited and represented as median and interquartile range for each of the variables calculated. A non-parametric Dunn’s test was used to compare the different conditions at 48 hours for *M. smegmatis* or 144 hours for *M. abscessus*. A significance level *a priori* was set at α = 0.05.

### Phylogenetic analysis of TPP pathway amino acid sequences in *M. abscessus*

A phylogenetic tree was constructed of a concatenated alignment of amino acid sequences of the five TPP synthesis pathway members for 143 clinical isolates of *M. abscessus* and *M. abscessus* ATCC19977. Homologs were identified using MMSeqs2 and phammseqs^47^ and subsequently aligned using ClustalO and Trimal^48^. A concatenated alignment was generated with a custom python script and the maximum likelihood phylogeny was generated using RAxML^49^.

