## Supplementary material for "*Mycobacterium* trehalose polyphleates are required for infection by therapeutically useful mycobacteriophages BPs and Muddy": All supplementary Information

**Supplementary Figure Legends**

**Figure S1. Plaque assays of phages on *M. abscessus* GD01 transposon insertion mutants**. Phages as shown on the left were spotted onto solid media with *M. abscessus* GD01 or transposon insertion mutant strains. Each row indicates a set strains tested together.

**Figure S2. Characterization of TPP-defective mutants. A.** Ziehl-Neelsen staining of *M. abscessus* TPP mutants. Cultures of GD22, GD22_RM, GD22_RM::C, GD180, GD180_RM and GD180_RM::C were fixed on glass slides and acid-fast staining was performed using the BD Carbolfuchsin kit, and observed microscopically at 60x magnification. **B.** MIC (in µg/mL) values of antibiotics determined in Cation-adjusted Mueller-Hinton Broth (CaMHB) at 30°C against *M. abscessus* clinical isolates (GD22 and GD180), resistant mutants (GD22_RM4 and GD180_RM2) and complemented strains (GD22_RM4::C and GD180_RM2::C) with CIP104536 (ATCC19977) as a control. Results from three independent experiments are shown.

**Figure S3. Infection of *M. smegmatis* TPP mutants by BPs and its derivatives A**. Growth curves of the different strains incubated with phage BPsΔ*33*HTH_HRM10 or BPsΔ*33*HTH_HRM10-mCherry at MOI 10 or without for 2 days in 7H9/OADC supplemented with 1 mM $CaCl_2$ at 37° C without agitation. Measurements were taken every 3 hours. Data shown are represented as median of three independents experiments made in triplicates ± interquartile range. Statistical analysis was done to compare the differences at 48 hours between each strain: ns: $P \geq 0.05$, * $P \leq 0.05$, *** $P \leq 0.001$, **** $P \leq 0.0001$. **B**. Representative microscope fields of *M. smegmatis* strains infected with the fluorophage BPsΔ*33*HTH_HRM10-mCherry (designated BPs mCherry) (MOI 10) for 2 hours at 37° C. Scale bars: 30 µm. **C**. Flow cytometry data plotted as a dot plot showing the percentage of bacilli infected with BPsΔ*33*HTH_HRM10-mCherry relative to the study population. **D**. Phage infection of *M. smegmatis* TPP mutants. Phages as shown on the left were ten-fold serially diluted and spotted onto solid media with *M. smegmatis* mc$^2$155, *M. smegmatis* Δ*papA3:*pLA155 or *M. smegmatis* Δ*papA3:*pKSW131. **E**. Plaquing of BPsΔ*33*HTH_HRM10 and the mCherry derivative on *M. smegmatis* strains defective in TPP production (left panels) and the complemented strains (right panels). Phage lysates were 10-fold serially diluted prior to spotting on bacterial lawns. Similar results were obtained at least three times and a representative experiment is shown.

**Figure S4. Gating strategy for flow cytometry analysis of fluorophage-infected bacteria.** Bacteria untreated or infected with non-fluorescent phages were used as controls. The first gate was plotted on SSC-A / FSC-A to exclude cellular debris. SSC-H / SSC-A was analysed to eliminate multiple cells. The gate for fluorescence was drawn thanks to controls to exclude non-fluorescent bacteria. The analysis was performed on approximately 300,000 bacteria using NovoExpress software. Here is shown an example of analysis for GD22 strain.

**Figure S5: Additional replicates of phage adsorption assays on *M. smegmatis* mc$^2$155 and Δ*pks*.** Replicates of phage adsorption assays for phages BPs_Δ33HTH_HRM10 **(A)**, phKSW1 **(B)** and ZoeJΔ43-45 **(C)** are shown. For BPs_Δ33HTH_HRM10, each individual assay showed reduced adsorption on Δ*pks* as compared to mc$^2$155. However, the measured adsorption rates varied between each replicate, such that observed differences were obfuscated when displaying the means; representative replicates for all phages are shown in Figure 5.

Table S1. Characterization of phage resistant *M. abscessus* GD01 transposon mutants

| Strain Name[1] | Isolated as resistant to[2] | # Insert[3] | Tn insertion coordinate[4] | Tn Dir[5] | GD01 Locus Tag[6] | ATCC19977 Gene[7] | Protein[8] | PCR ver.[9] | Plaque assay[10] |
|---|---|---|---|---|---|---|---|---|---|
| GD01Tn_Muddy_RM9 | Muddy | ≥1 | 895811 | Rev | EXM25_04535 | MAB_0935c | FadD23 | Y | Y |
| GD01Tn_BPsHRM10_RM9 | BPs_HRM10 | ≥1 | 896103 | Fwd | EXM25_04535 | MAB_0935c | FadD23 | ND | ND |
| GD01Tn_BPsHRM10_RM15 | BPs_HRM10 | ≥1 | 896250 | Rev | EXM25_04535 | MAB_0935c | FadD23 | ND | ND |
| GD01Tn_Muddy_RM14 | Muddy | ≥2 | #1 896251<br>#2 2068078 | Fwd<br>Fwd | EXM25_04535<br>EXM25_10305 | MAB_0935c<br>MAB_2130 | FadD23<br>HAD | Y<br>Y | Y |
| GD01Tn_Muddy_RM11[11] | Muddy | ≥1 | 896372 | Rev | EXM25_04535 | MAB_0935c | FadD23 | ND | ND |
| GD01Tn_Muddy_RM15[11] | Muddy | ≥1 | 896372 | Rev | EXM25_04535 | MAB_0935c | FadD23 | ND | ND |
| GD01Tn_Muddy_RM13 | Muddy | ≥1 | 896513 | Rev | EXM25_04535 | MAB_0935c | FadD23 | ND | ND |
| GD01Tn_Muddy_RM7 | Muddy | ≥1 | 896840 | Rev | EXM25_04535 | MAB_0935c | FadD23 | ND | ND |
| GD01Tn_Muddy_RM12 | Muddy | ≥1 | 896990 | Rev | EXM25_04535 | MAB_0935c | FadD23 | Y | Y |
| GD0Tn_BPsHRM10_RM1 | BPs_HRM10 | ≥1 | 897275 | Rev | EXM25_04535 | MAB_0935c | FadD23 | Y | Y |
| GD01Tn_BPsHRM10_RM6 | BPs_HRM10 | ≥1 | 897828 | Rev | EXM25_04540 | MAB_0936c | PE | Y | Y |
| GD01Tn_BPsHRM10_RM3 | BPs_HRM10 | ≥2 | #1 898266<br>#2 1702803 | Fwd<br>Rev | EXM25_04540<br>EXM25_08515 | MAB_0936c<br>MAB_1681 | PE<br>Hyp | Y<br>Y | Y |
| GD01Tn_BPsHRM10_RM8 | BPs_HRM10 | | #1 898475<br>#2 2710182 | Fwd<br>Rev | EXM25_04540<br>EXM25_13435 | MAB_0936c<br>MAB_2752 | PE<br>ABC | ND<br>ND | ND |
| GD01Tn_Muddy_RM20 | Muddy | | 898646 | Rev | EXM25_04540 | MAB_0936c | PE | Y | Y |
| GD01Tn_BPsHRM10_RM18 | BPs_HRM10 | ≥2 | #1 898656<br>#2 50130 | Rev<br>Fwd | EXM25_04540<br>EXM25_00255 | MAB_0936c<br>MAB_0457 | PE<br>Hyp | Y<br>Y | Y |
| GD01Tn_BPsHRM10_RM20 | BPs_HRM10 | ≥3 | #1 899155<br>#2 2314064<br>#3 4430814 | Fwd<br>Fwd<br>Fwd | EXM25_04545<br>EXM25_11400<br>Inter EXM25_22125-22130 | MAB_0937c<br>MAB_2352<br>MAB_4446-4447 | MmpL10<br>DNA glyc<br>Hyp/ABC | ND<br>ND<br>ND | ND |
| GD01Tn_BPsHRM10_RM11 | BPs_HRM10 | ≥1 | 899212 | Fwd | EXM25_04545 | MAB_0937c | MmpL10 | Y | Y |
| GD01Tn_BPsHRM10_RM14 | BPs_HRM10 | ≥2 | #1 900526<br>#2 3419195 | Rev<br>Rev | EXM25_04545<br>EXM25_17025 | MAB_0937c<br>MAB_3426c | MmpL10<br>SDR | Y<br>Y | Y |
| GD01Tn_BPsHRM10_RM19 | BPs_HRM10 | ≥2 | #1 901936<br>#2 2004769 | Fwd<br>Rev | EXM25_04550<br>EXM25_10050 | MAB_0938c<br>MAB_2081 | PapA3<br>Acyl-CoA | Y<br>Y | Y |
| GD01Tn_BPsHRM10_RM2 | BPs_HRM10 | ≥1 | 901937 | Rev | EXM25_04550 | MAB_0938c | PapA3 | ND | ND |
| GD01Tn_Muddy_RM17 | Muddy | ≥1 | 902798 | Fwd | EXM25_04550 | MAB_0938c | PapA3 | Y | Y |
| GD01Tn_BPsHRM10_RM10 | BPs_HRM10 | ≥1 | 902799 | Rev | EXM25_04550 | MAB_0938c | PapA3 | Y | Y |
| GD01Tn_Muddy_RM1[11] | Muddy | ≥2 | 902799<br>1708979 | Rev<br>Fwd | EXM25_04550<br>EXM25_08540 | MAB_0938c<br>MAB_1686 | PapA3<br>Hyp | ND<br>ND | ND |
| GD01Tn_Muddy_RM3[11] | Muddy | ≥2 | 902799<br>1708979 | Rev<br>Fwd | EXM25_04550<br>EXM25_08540 | MAB_0938c<br>MAB_1686 | PapA3<br>Hyp | ND<br>ND | ND |
| GD01Tn_Muddy_RM6 | Muddy | ≥2 | #1 902802<br>#2 1799970 | Rev<br>Fwd | EXM25_04550<br>EXM25_09000 | MAB_0938c<br>MAB_1882c | PapA3<br>DoxX | Y<br>Y | Y |
| GD01Tn_Muddy_RM19 | Muddy | | 903001 | Rev | EXM25_04550 | MAB_0938c | PapA3 | Y | Y |
| GD01Tn_BPsHRM10_RM4 | BPs_HRM10 | ≥2 | #1 903920<br>#2 3438375 | Fwd<br>Fwd | EXM25_04555<br>EXM25_17140 | MAB_0939<br>MAB_3449c | Pks<br>MFS | Y<br>Y | Y |
| GD01Tn_BPsHRM10_RM5 | BPs_HRM10 | ≥2 | #1 905614<br>#2 3575341 | Fwd<br>Fwd | EXM25_04555<br>EXM25_17815 | MAB_0939<br>MAB_3593 | Pks<br>MgtC | Y<br>Y | Y |
| GD01Tn_BPsHRM10_RM17 | BPs_HRM10 | ≥2 | #1 911720<br>2# 3319297 | Rev<br>Rev | EXM25_04555<br>EXM25_16510 | MAB_0939<br>MAB_3332c | Pks<br>DoxX | Y<br>Y | Y |
| GD01Tn_BPsHRM10_RM13 | BPs_HRM10 | ≥2 | #1 914389<br>#2 934573 | Fwd<br>Rev | EXM25_04555<br>EXM25_04650 | MAB_0939<br>MAB_0958c | Pks<br>NAD(P) | Y<br>Y | Y |
| GD01Tn_BPsHRM10_RM7 | BPs_HRM10 | ≥2 | #1 914477 | Rev | EXM25_04555 | MAB_0939 | Pks | Y | Y |

| | | | | | | | | | |
|---|---|---|---|---|---|---|---|---|---|
| | | | #2 3546108 | Rev | Inter EXM25_17670-17675 | MAB_3538-3539 | Kinase/ WhiB | Y | |
| GD01Tn_phKSW1_RM1[11] | phKSW1 | Prob 1 | 1711653 | Fwd | EXM25_08560 | MAB_1690 | ABC | Y | Y |
| GD01Tn_phKSW1_RM4[11] | phKSW1 | Prob 1 | 1711653 | Fwd | EXM25_08560 | MAB_1690 | ABC | Y | Y |
| GD01Tn_phKSW1_RM2 | phKSW1 | Prob 1 | 1169901 | Fwd | Inter EXM25_05825-05830 | None – MAB_1175 | DUF4145 | Y | Y |
| GD01Tn_phKSW1_RM7[11] | phKSW1 | ≥2 | #1 1169901 | Fwd | Inter EXM25_05825-05830 | None – MAB_1175 | DUF4145 | Y | Y |
| | | | #2 2201133 | Fwd | EXM25_10900 | MAB_2245 | | Y | |
| GD01Tn_phKSW1_RM8[11] | phKSW1 | ≥2 | #1 1169901 | Fwd | Inter EXM25_05825-05830 | None – MAB_1175 | DUF4145 | Y | Y |
| | | | #2 2201133 | Fwd | EXM25_10900 | MAB_2245 | | Y | |
| GD01Tn_phKSW1_RM5 | phKSW1 | ≥2 | #1 1169902 | Rev | Inter EXM25_05825-05830 | None – MAB_1175 | DUF4145 | Y | Y |
| | | | #2 4426058 | Rev | EXM25_22105 | MAB_4441 | | Y | |
| GD01Tn_phKSW1_RM6 | phKSW1 | ≥2 | #1 3688289 | Fwd | EXM25_18375 | MAB_3698 | ATP | Y | Y |
| | | | #2 428992 | Fwd | EXM25_02155 | MAB_0399c | RecB | Y | |
| GD01Tn_phKSW1_RM10 | phKSW1 | ≥2 | #1 1708979 | Fwd | EXM25_08540 | MAB_1686 | Hyp | Y | Y |
| | | | #2 902799 | Rev | EXM25_04550 | MAB_0938c | PapA3 | Y | |
| GD01Tn_phKSW1_RM11a | phKSW1 | ≥1 | 3371296 | Rev | EXM25_16760 | MAB_3381c | Mtf | Y | Y |
| GD01Tn_MuddyREM1_RM2 | Muddy_REM1 | ≥1 | 1195603 | Rev | EXM25_05970 | MAB_1201c | GreA | Y | Y |
| GD01Tn_MuddyREM1_RM3 | Muddy_REM1 | Prob 1 | 3850846 | Rev | EXM25_19265 | MAB_3868c | RpoC | Y | Y |

[1] Strain names include three parameters separated by underscores: i.e. Parent strain_phage used for selection_mutant number.
[2] The strain was isolated from a solid agar plate seeded with phages as indicated.
[3]The likely number of Tn insertions in each strain: Prob 1 indicates that only one insertion was identified and mapped, and that other insertions are unlikely but cannot be excluded; ≥1 indicates that we have mapped one insertion, but there are one or more additional insertions that have not been mapped; ≥2 indicates that two insertions were mapped, but we cannot exclude that there are one or more additional insertions.
[4]Coordinates of the transposon insertions mapped in *M. abscessus* GD01 (accession number CP035923.1). Where more than one Tn has been mapped, they are listed as #1 and #2, and #2 is shown in grey shading. Strains with similar coordinates are listed together.
[5]Orientation of the Tn insertion, Forward (Fwd) or reverse (Rev) compared to the genome.
[6]The locus tag of the strain GD01 gene with the Tn insertion is shown. Intergenic insertions show the two flanking locus tags.
[7]The gene name of the *M. abscessus* ATCC 19977 (NC_010397.1) homolog(s) of the interrupted genes is shown. For intergenic insertions, flanking genes are shown.
[8]Protein names or predicted function of genes with Tn insertions are shown, or functions of flanking genes for intergenic insertions.
[9]The location of the transposon insertion was confirmed by a second PCR with primers designed to flank the insertion site. Y, Yes; ND, Not determined.
[10]The strain was streaked out to remove phage, grown in liquid media and tested for phage resistance by plaque assay. Y, Yes; ND, Not determined.
[11]In four instances, two strains isolated against the same phage were found to have identical transposon insertion sites and are likely siblings. GD01Tn_phKSW1_RM1 and GD01Tn_phKSW1_RM4 are likely siblings, as are GD01Tn_phKSW1_RM7 and GD01Tn_phKSW1_RM8. GD01Tn_phKSW1_RM2, GD01Tn_phKSW1_RM5, and GD01Tn_phKSW1_RM7 have insertions in the same location (coordinate 1169901) but are not siblings as they have Tn insertions in different orientations or different secondary insertions.

Table S2. Phage BPs and Muddy mutants escaping TPP-loss mediated resistance

| Phage[1] | Parent[2] | Amino Acid Substitution[3] | Isolated on[4] | Reference |
|---|---|---|---|---|
| phKSW2 | BPsΔ*33*HTH_HRM10 | gp22 L462R | *M. ab.* GD01Tn BPs_HRM10_RM6 | This work |
| phKSW3 | BPsΔ*33*HTH_HRM10 | gp22 A306V | *M. ab.* GD01Tn BPs_HRM10_RM6 | This work |
| phKSW4 (Prob. a sib of phKSW2) | BPsΔ*33*HTH_HRM10 | gp22 L462R | *M. ab.* GD01Tn BPs_HRM10_RM11 | This work |
| phKSW5 | BPsΔ*33*HTH_HRM10 | gp22 A604E | *M. ab.* GD01Tn BPs_HRM10_RM11 | This work |
| BPs_REM1 | BPsΔ*33*HTH_HRM10 | gp22 L462R; G780R | *M. ab.* GD180_RM2 | This work |
| phKSW1 | BPsΔ*33*HTH_HRM10 | gp22 A604E | *M. smegmatis* Δ*MSMEG_5439* | This work |
| Muddy_REM1 | Muddy | gp24 E680K | *M. ab.* GD180_RM2 | This work |
| Muddy_HRM$^{N0157}$-1 | Muddy | gp24 G487W | *M. tuberculosis* N0157 | [1] |
| Muddy_HRM$^{N0157}$-2 | Muddy | gp24 T608A | *M. tuberculosis* N0157 | [1] |
| Muddy_HRM$^{N0052}$-1 | Muddy | gp24 E680K | *M. tuberculosis* N0052 | [1] |

[1] Name of a phage able to form more clear plaques on TPP synthesis pathway mutants
[2] Parent phage from which the mutant was isolated
[3] Amino acid substitutions identified in mutant phages
[4] The *M. abscessus* or *M. tuberculosis* strain on which the mutant phage was isolated

Table S3. *M. abscessus* mutants spontaneously resistant to phage BPs derivatives

| Resistant mutant[1] | Parent[2] | Isolated as resistant to[3] | Mutated gene[4] | Amino Acid Substitution[5] | Reference |
|---|---|---|---|---|---|
| GD17_RM1 | *M. ab.* GD17 | BPsHRM$^{GD03}$ | *pks* | G210V | [2] |
| GD22_RM4 | *M. ab.* GD22 | BPsΔ*33*HTH_ HRM10 and Itos | *pks* | W2389fsX2406 | [2] |
| GD38_RM2 | *M. ab.* GD38 | BPsΔ*33*HTH_ HRM10 | *pks* | M2115fsX2118 | This work |
| GD59_RM1 | *M. ab.* GD59 | BPsΔ*33*HTH_ HRM10 and Itos | *pks* | D327Y | This work |
| GD180_RM2 | *M. ab.* GD180 | BPsΔ*33*HTH_ HRM10 | *mmpL10* | S688fsX704 | This work |

[1] Resistant mutants were named after the parent strain and the number of the resistant mutant (RM) isolated
[2] Parent strain of *M. abscessus* from which the resistant mutant was isolated upon phage infection
[3] RMs were isolated by infecting cell cultures at a high MOI and plating infections on solid media to isolate survivors
[4] The entire genomes of the resistant mutants were sequenced and compared to the genomes of the parents to identify mutated genes conferring phage resistance
[5] The amino acid substitution in the protein product resulting from the gene mutation

Table S4. List of strains used in this study

| Name | Description / genotype | Reference |
|---|---|---|
| *M. smegmatis* (WT) | *M. smegmatis*, strain mc²155 | 3 |
| Δ*pks* (PMM284) | *M. smegmatis* mc²155 Δ*pks::res* | 4 |
| Δ*mmpL10* (PMM223) | *M. smegmatis* mc²155 Δ*mmpL10::res* | 4 |
| Δ*pE* (PMM229) | *M. smegmatis* mc²155 Δ*pE::res* | 4 |
| Δ*pks*::C (PMM284/pMV*pks*) | *M. smegmatis* mc²155 Δ*pks::res* complemented with pMV*pks*, $Hyg^R$ | 4 |
| Δ*mmpL10*::C (PMM223/pMV*mmpL10*) | *M. smegmatis* mc²155 Δ*mmpL10::res* complemented with pMV*mmpL10*, $Hyg^R$ | 4 |
| Δ*pE*::C (PMM229/pMV*pE*) | *M. smegmatis* mc²155 Δ*pE::res* complemented with pMV*pE*, $Hyg^R$ | 4 |
| GD01 | *M. abscessus subsp. massiliense,* R morphotype | 5 |
| GD17 | *M. abscessus subsp. abscessus,* R morphotype | 2 |
| GD17_RM1 | *M. abscessus subsp. abscessus,* R morphotype | 2 |
| GD22 | *M. abscessus subsp abscessus*, R morphotype | 2 |
| GD22_RM4 | *M. abscessus subsp. abscessus* spontaneous resistant mutant, R morphotype | 2 |
| GD22_RM4::C | *M. abscessus* GD22_RM4 complemented with pMV*pks_mWasabi*, $Hyg^R$ | This study |
| GD38 | *M. abscessus subsp. abscessus,* R morphotype | 2 |
| GD59_RM1 | *M. abscessus subsp. abscessus,* R morphotype | 2 |
| GD59 | *M. abscessus subsp. abscessus,* R morphotype | 2 |
| GD180 | *M. abscessus subsp. abscessus*, R morphotype | This study |
| GD180_RM2 | *M. abscessus subsp. abscessus,* spontaneous resistant mutant, R morphotype | This study |
| GD180_RM2::C | *M. abscessus* GD180_RM2 complemented with pMV*mmpL10_mWasabi*, $Hyg^R$ | This study |
| Δ*fadD23* | *M. smegmatis* mc²155 Δ*fadD23::ZeoR* | 6 |
| Δ*fadD23*:pKSW131 | *M. smegmatis* mc²155 Δ*fadD23* complemented with pKSW131 | This study |
| Δ*papA3* | *M. smegmatis* mc²155 Δ*papA3::ZeoR* | 6 |
| GD273 | *M. abscessus* subsp. massiliense | This study |
| GD286 | *M. abscessus subsp. massiliense* | This study |
| Δ*papA3 :pLA155* | *M. smegmatis* mc²155 Δ*fadD23* complemented with pLA155 | This study |
| Δ*papA3 :pKSW131* | *M. smegmatis* mc²155 Δ*papA3* complemented with pKSW131 | This study |

Table S5. Plasmids and primers used in this study.

| Name | Plasmid/Primer | Description |
|---|---|---|
| pMV*pks_mWasabi* | Plasmid | pMV361eH containing *pks* under the control of its own promoter[1] and *mWasabi* sequence under the control of the constitutive *Pleft**promoter (addgene plasmid :169409) |
| pMV*mmpL10_mWasabi* | Plasmid | pMV361eH containing *mmpL10* under the control of the *PblaF** promoter[1] and *mWasabi* sequence under the control of the constitutive *Pleft**promoter (addgene plasmid :169409) |
| pLA155 | Plasmid | pMH94 (L5 integrase) where the Kanamycin resistance cassette has been replaced with a streptomycin resistance cassette |
| pKSW131 | Plasmid | pLA155 containing *papA3*, *mmpL10*, *pE* and *fadD23* from *M. abscessus* GD01 under control of the native promoter |
| pKSW134 | Plasmid | pCCK39 (L5 integrase, streptomycin resistance cassette ) containing *fadD23* from *M. abscessus* GD01 under control of a tet-inducible promoter. |
| Inf_*Pleft**_*mWasabi*_pMV (KpnI) (F) | Primer | 5'-CCACTGCGATCCCCGGGTACTGATGCCTGGCAGTCGATCGT |
| Inf_*Pleft**_*mWasabi*_pMV (KpnI) (R) | Primer | 5'- CGTCGCCGAGGGCTTGGTACGGCCGCGGTACCAGATCTT |
| TPP_pMH94_Fwd | Primer | TTGTAAAACGACGGCCAGTGAATTCTTGTGGCCTCCTTGCGTC |
| TPP_pMH94_Rev | Primer | ATCCCCGGGTACCGAGCTCGAATTCACCGAGAGGCTAGGCGAC |
| FadD_pCCK39_Fwd | Primer | GATTCGCCGCCCGAAATCACAGCGTGACTTGGACAAAACTATGACACC |
| FadD-pCCK39_Rev | Primer | GCGTTTAAACCTGCAGGCACCTAGGCGACGCCCACCGG |

A

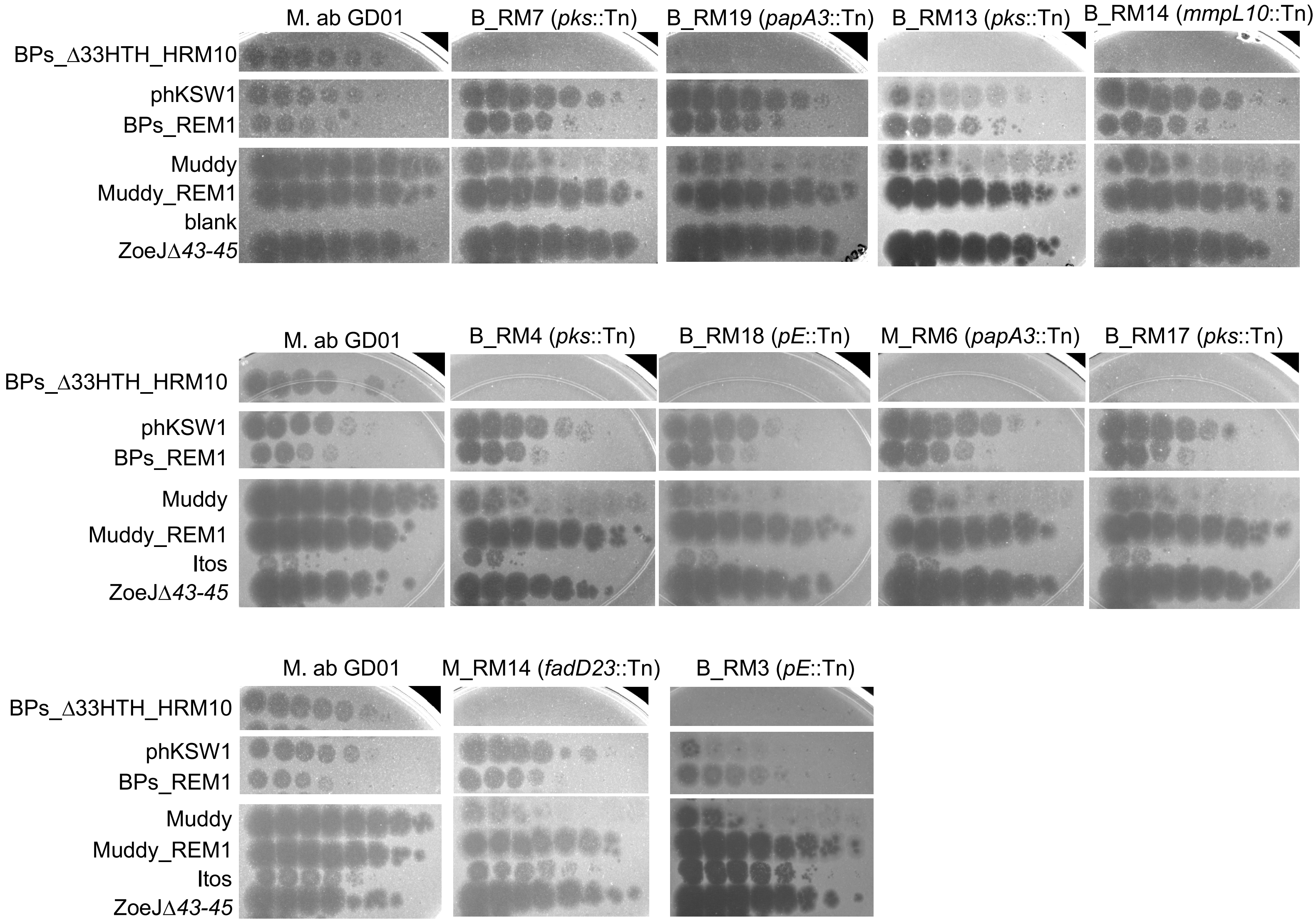
M. ab GD01
B_RM7 (pks::Tn)
B_RM19 (papA3::Tn)
B_RM13 (pks::Tn)
B_RM14 (mmpL10::Tn)
BPs_Δ33HTH_HRM10
phKSW1
BPs_REM1
Muddy
Muddy_REM1
blank
ZoeJΔ43-45
M. ab GD01
B_RM4 (pks::Tn)
B_RM18 (pE::Tn)
M_RM6 (papA3::Tn)
B_RM17 (pks::Tn)
BPs_Δ33HTH_HRM10
phKSW1
BPs_REM1
Muddy
Muddy_REM1
Itos
ZoeJΔ43-45
M. ab GD01
M_RM14 (fadD23::Tn)
B_RM3 (pE::Tn)
BPs_Δ33HTH_HRM10
phKSW1
BPs_REM1
Muddy
Muddy_REM1
Itos
ZoeJΔ43-45

Figure S1

A

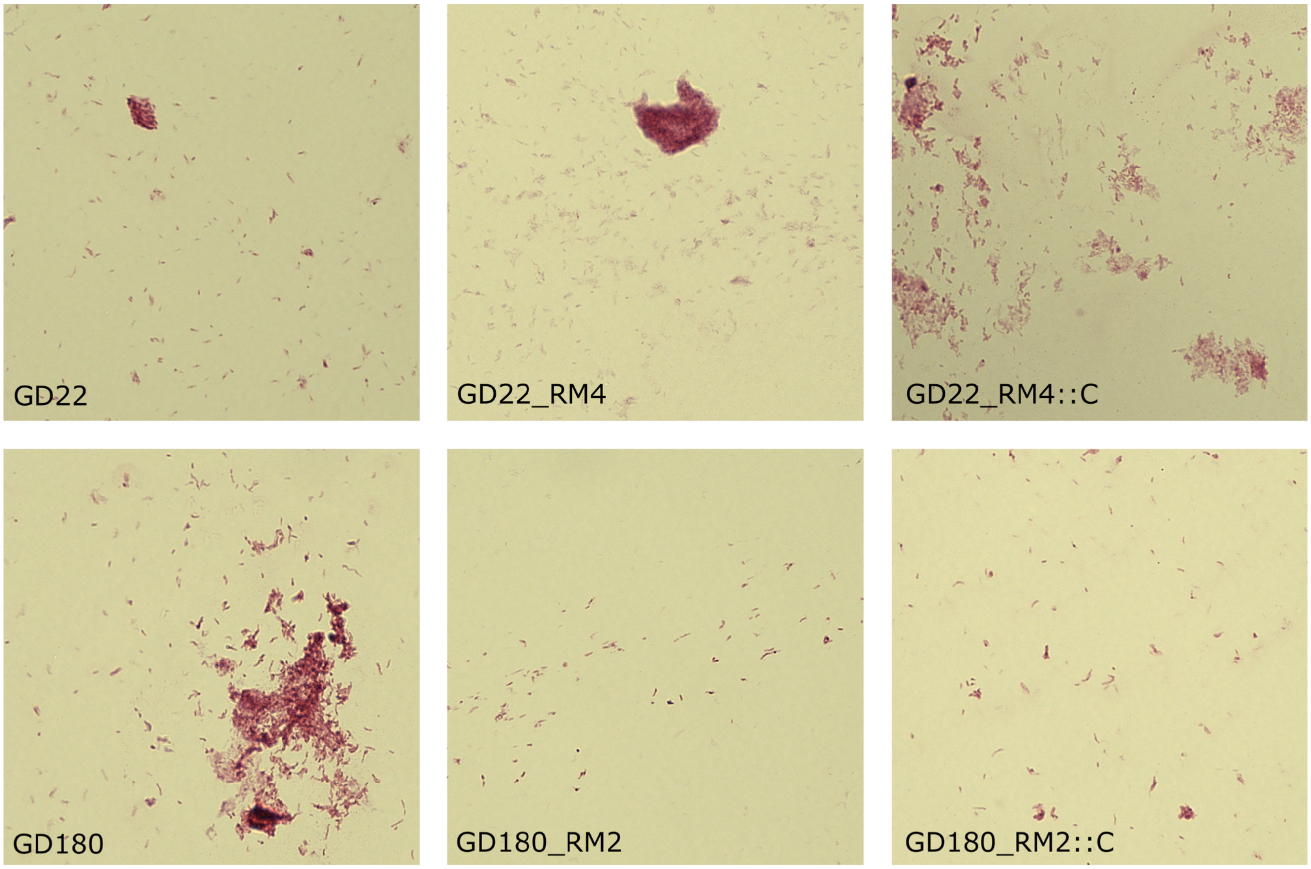


B

| Strains | | MIC (µg/mL) [a] | | | | | | | | |
|---|---|---|---|---|---|---|---|---|---|---|
| | | AMK | IPM | BDQ | RFB | CFZ | LNZ | CFX | ZEO | AU1235 |
| **GD22** | Exp 1 | 25 | 25-50 | 0.098 | 12.5 | 3.125 | 12.5 | 50-100 | 12.5 | 0.39 |
| | Exp 2 | 50 | 50 | 0.098 | 25 | 1.56 | 25 | 100 | 12.5 | 0.195 |
| | Exp 3 | 100 | 50 | 0.098 | 25 | 0.39 | 6.25 | 100 | 6.25 | 0.39 |
| **GD22_RM4** | Exp 1 | 25 | 25 | 0.195 | 12.5 | 1.56 | 6.25 | 50 | 6.25 | 0.78 |
| | Exp 2 | 50 | 25 | 0.098 | 12.5 | 1.56 | 12.5 | 50 | 6.25 | 0.195 |
| | Exp 3 | 50 | 100 | 0.098 | 25 | 3.25 | 12.5 | 50 | 12.5 | 0.195 |
| **GD22_RM4 :: C** | Exp 1 | 50 | 12.5 | 0.098 | 12.5 | 1.56 | 12.5 | 50 | 6.25 | 0.39 |
| | Exp 2 | 50 | 25 | 0.098 | 12.5 | 0.78 | 12.5 | 50 | 6.25 | 0.195 |
| | Exp 3 | 50 | 25-50 | 0.19 | 25 | 1.56 | 6.25 | 50 | 6.25 | 0.195 |
| **GD180** | Exp 1 | 25 | 25 | 0.098 | 1.56 | ND | 12.5 | 25 | >200 | 0.048 |
| | Exp 2 | 25 | 50 | 0.048 | 6.25 | 0.78 | 6.25 | 50 | >200 | <0.048 |
| | Exp 3 | 50 | 25 | 0.024 | 12.5 | 0.78 | 6.25 | 25 | >200 | 0.19 |
| **GD180_RM2** | Exp 1 | 25 | 12.5 | 0.024 | 3.125 | 1.56 | 6.25 | 25 | >200 | 0.39 |
| | Exp 2 | 50 | 25 | 0.098 | 3.125 | 1.56 | 1.56 | 25 | >200 | <0.048 |
| | Exp 3 | 50 | 12.5 | 0.048 | 6.25 | 0.78 | 1.56 | 25 | >200 | 0.19 |
| **GD180_RM2 :: C** | Exp 1 | 50 | 25 | 0.098 | 3.125 | 0.78 | 3.125 | 25 | >200 | 0.098 |
| | Exp 2 | 25 | 50 | 0.024 | 3.125 | 0.78 | 3.125 | 50 | >200 | <0.048 |
| | Exp 3 | 50 | 25 | 0.024 | 3.125 | 0.78 | 3.125 | 25 | >200 | 0.098 |
| **CIP104536 (R)** | Exp 1 | 50 | 50 | 0.048 | 12.5-25 | 1.56 | 25 | 100 | 50 | 0.39 |
| | Exp 2 | 50 | 25 | 0.098 | 25 | 0.78 | 12.5 | 50 | 25 | 0.78 |

[a] MICs (µg/ml) were determined following the CLSI guidelines.

AMK, amikacin; IPM, imipenem; BDQ, bedaquiline; RFB,rifabutin; CFZ, clofazimine; LNZ, linezolid; CFX, cefoxitin; ZEO, zeocin; AU1235, MmpL3 inhibitor

Figure S2

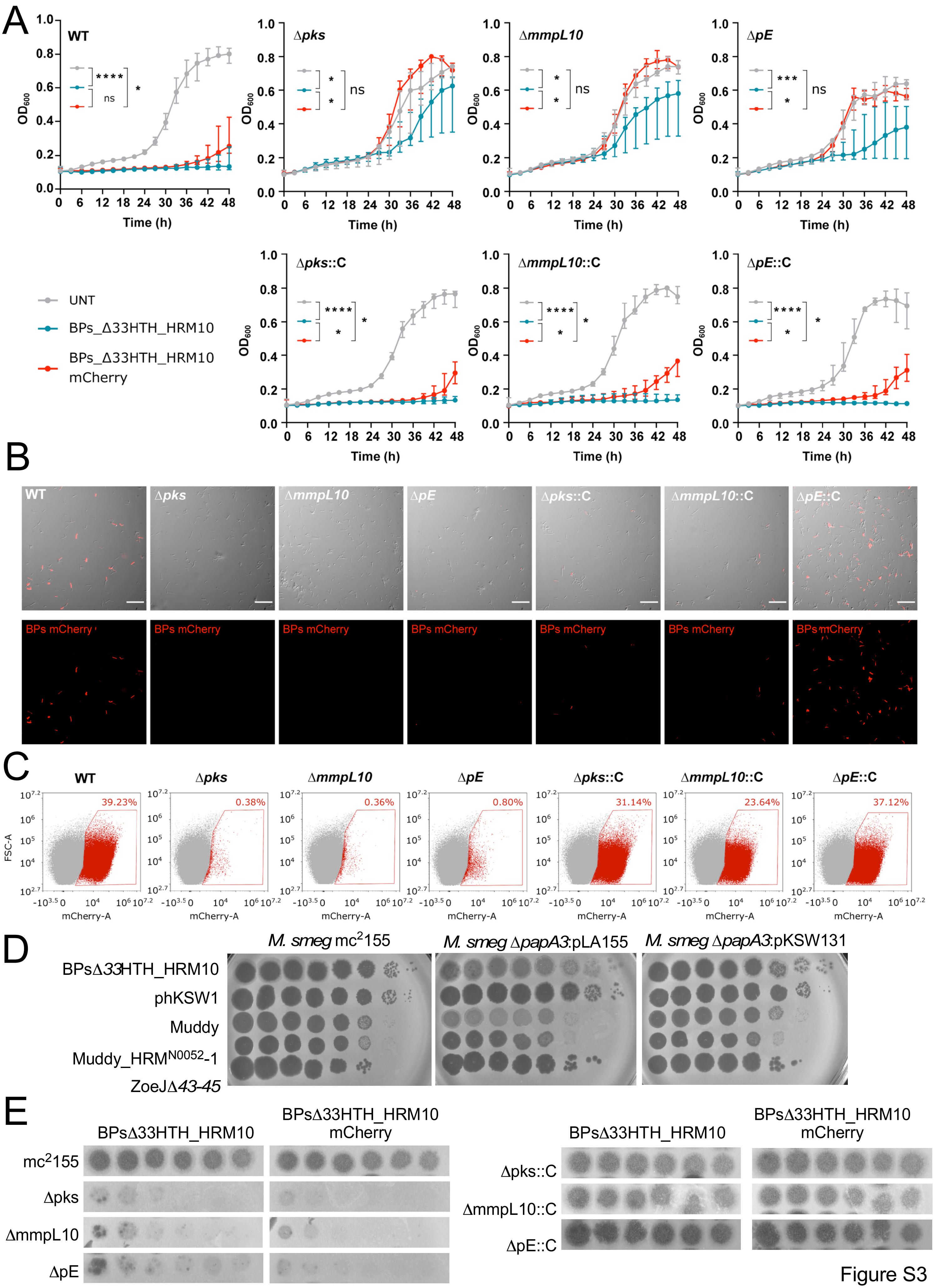
A
WT
Δpks
ΔmmpL10
ΔpE
Δpks::C
ΔmmpL10::C
ΔpE::C
UNT
BPs_Δ33HTH_HRM10
BPs_Δ33HTH_HRM10 mCherry
Time (h)
OD600
B
BPs mCherry
C
39.23%
0.38%
0.36%
0.80%
31.14%
23.64%
37.12%
FSC-A
mCherry-A
D
M. smeg mc2155
M. smeg ΔpapA3:pLA155
M. smeg ΔpapA3:pKSW131
BPsΔ33HTH_HRM10
phKSW1
Muddy
Muddy_HRMN0052-1
ZoeJΔ43-45
E
BPsΔ33HTH_HRM10
BPsΔ33HTH_HRM10 mCherry
mc2155
Δpks
ΔmmpL10
ΔpE
Δpks::C
ΔmmpL10::C
ΔpE::C

Figure S3

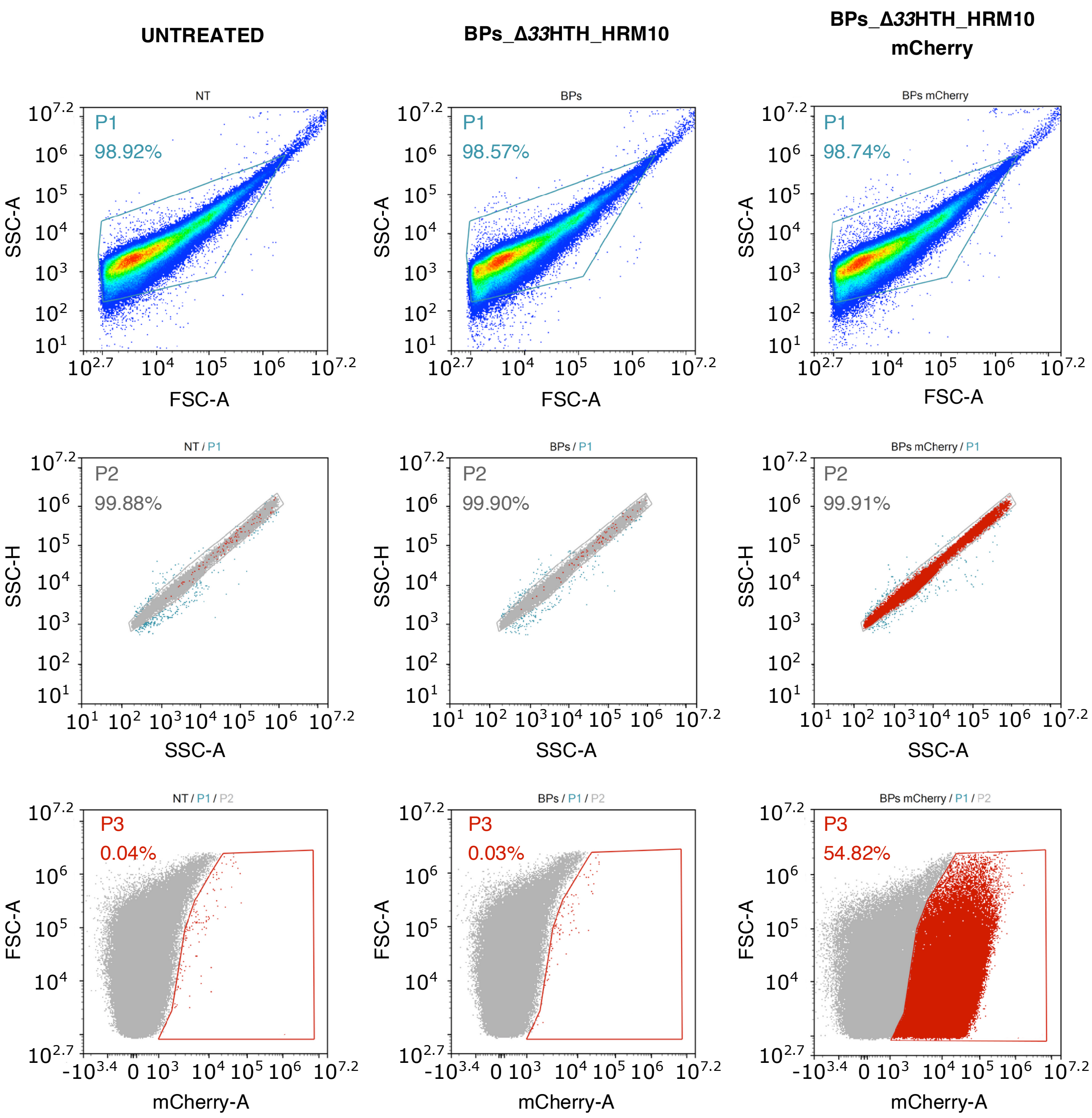

UNTREATED
BPs_Δ33HTH_HRM10
BPs_Δ33HTH_HRM10
mCherry
NT
BPs
BPs mCherry
P1
98.92%
98.57%
98.74%
SSC-A
FSC-A
NT / P1
BPs / P1
BPs mCherry / P1
P2
99.88%
99.90%
99.91%
SSC-H
SSC-A
NT / P1 / P2
BPs / P1 / P2
BPs mCherry / P1 / P2
P3
0.04%
0.03%
54.82%
FSC-A
mCherry-A


Figure S4

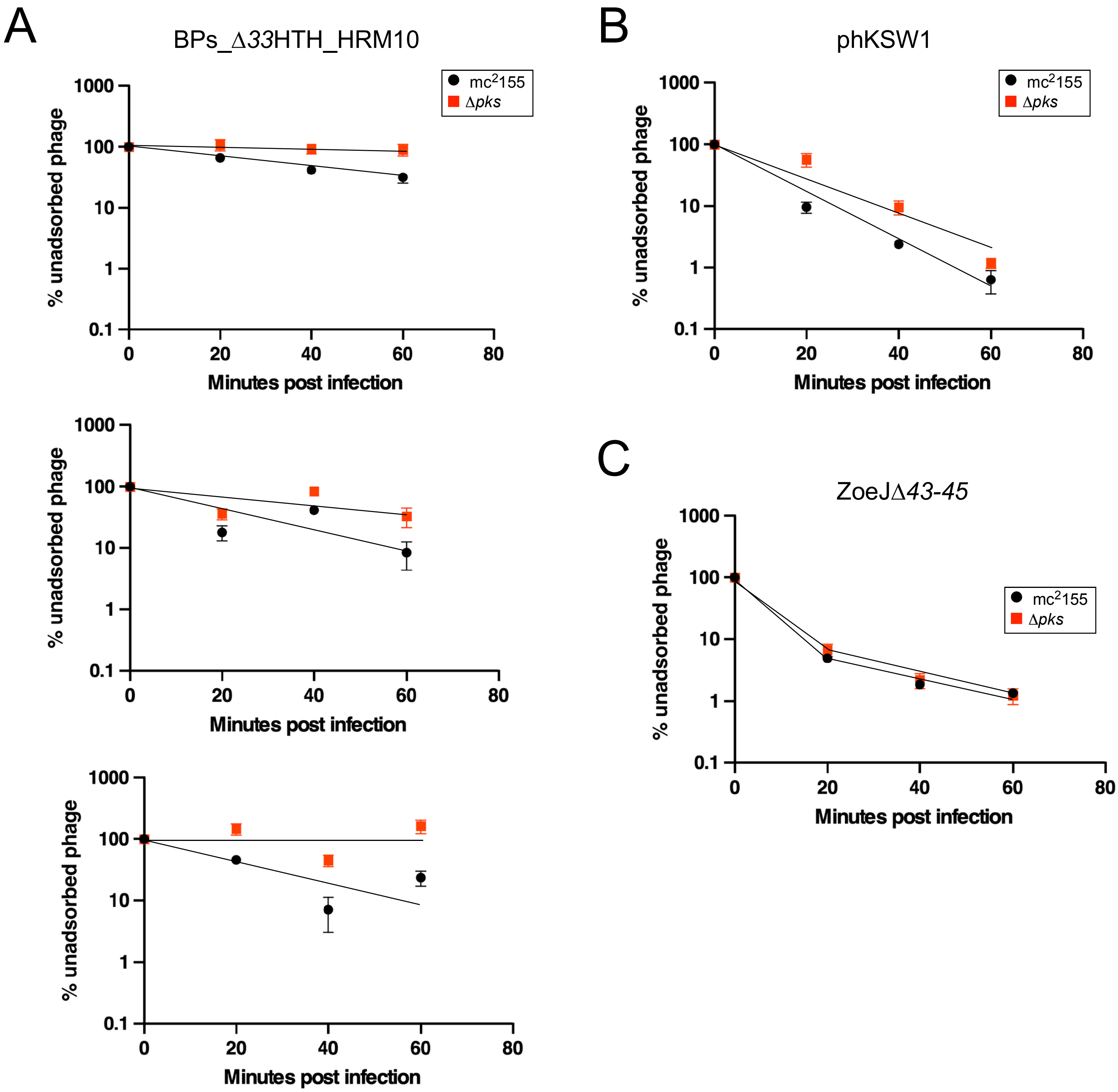
A
BPs_Δ33HTH_HRM10
B
phKSW1
C
ZoeJΔ43-45
mc2155
Δpks
% unadsorbed phage
Minutes post infection
1000
100
10
1
0.1
0
20
40
60
80

Figure S5
